# The Oviposition Inhibitory Neuron is a potential hub of multi-circuit integration in the *Drosophila* brain

**DOI:** 10.1101/2024.10.25.620362

**Authors:** Rhessa A. Weber Langstaff, Pranjal Srivastava, Alexander B. Kunin, Gabrielle J. Gutierrez

## Abstract

Understanding how neural circuits integrate sensory and state information to support context-dependent behavior is a central challenge in neuroscience. Oviposition is a complex process during which a fruit fly integrates context and sensory information to choose an optimal location to lay her eggs. The circuit that controls oviposition is known, but how the oviposition circuit integrates multiple sensory modalities and internal states is not. Using the Hemibrain connectome, we identified the Oviposition Inhibitory Neuron (oviIN) as a key hub in the oviposition circuit and analyzed its inputs to uncover potential parallel pathways that may be responsible for computations related to sensory integration and decision-making. We applied a network analysis to the subconnectome of inputs to the oviIN to identify clusters of interconnected neurons – many of which are uncharacterized cell types. Our findings indicate that the inputs to oviIN form multiple parallel pathways through the unstructured neuropils of the Superior Protocerebrum, a region implicated in context-dependent processing.

**Significance Statement:** The recent advent of the *Drosophila* connectome enables researchers to probe the connectivity of uncharacterized cell types in the parts of the fruit fly brain that are responsible for cognitive-level computations. Our study analyzed the connectivity of the oviposition circuit which controls a complex behavior that depends on sensory and context integration and decision-making computations. Using graph theoretic and computational methods, we found that the sole inhibitory neuron in the circuit is a hub that integrates information from multiple clusters of uncharacterized neurons with potentially novel functions. Our work presents a new and timely perspective by demonstrating how new targets for study can be identified from the vast trove of uncharacterized neurons and cell types in the connectome.

## Introduction

Identifying the circuits responsible for cognitive-level processing within the vast network of the fruit fly brain presents a significant challenge (***O’Leary and Marder, 2014***). The recent advent of partial and full-brain connectomes provides a powerful resource for this investigation (***Scheffer et al., 2020***; ***Dorkenwald et al., 2024***; ***Schlegel et al., 2024***). Publicly accessible datasets provide the most comprehensive map of the connectivity in the fruit fly brain currently available, although there is some variability between connectomes (***Lin et al., 2024***).

Oviposition is a prime example of a behavior that relies on cognitive-level processing (***Cury et al., 2019***; ***Yang et al., 2008***; ***Churchill et al., 2021***; ***Vijayan et al., 2023***; ***Cury and Axel, 2023***; ***Yang et al., 2015***). Following mating, the female fly assesses a variety of factors such as the firmness (***Zhang et al., 2020***), taste (***Vijayan et al., 2022***; ***Joseph et al., 2009***; ***Yang et al., 2008***), and spatial attributes of a substrate (***Schwartz et al., 2012***), before deciding whether to deposit an egg. While sampling and evaluating multiple sensory inputs, the fly computes a relative value for each encountered substrate (***Vijayan et al., 2023***; ***Yang et al., 2008***, ***2015***; ***Azanchi et al., 2013***). The fly considers social contexts and physical environment to optimize survival of the egg (***Rockwell and Grossfield, 1978***; ***Shelly, 1999***; ***Schwartz et al., 2012***; ***Bailly et al., 2023***). Computations such as these contribute to the complex decision-making process inherent in oviposition.

The circuit responsible for commanding oviposition and the cell types involved are known (***Wang et al., 2020***). Within each hemisphere, a set of Oviposition Descending Neurons (oviDN) are directly downstream of a single Oviposition Excitatory Neuron (oviEN) and Oviposition Inhibitory Neuron (oviIN) (***Figure 1***A). Mating status is conveyed to the oviDN via the oviIN by way of a subset of pC1 neurons. The pC1 cluster of neurons is a sexually dimorphic cell type that promotes persistent aggressive behaviors (***Deutsch et al., 2020***; ***Schretter et al., 2020***) and sexual receptivity (***Zhou et al., 2014***). After a mating experience, sex peptide inhibits the Sex Peptide Sensory Neurons (SPSN) and Sex peptide Abdominal Ganglia (SAG) neurons (***Wang et al., 2020***; ***Feng et al., 2014***; ***Laturney et al., 2023***). The pC1a and pC1b neurons downstream of SPSN and SAG are silenced by the reduced input which in turn silences the oviIN. The result is the disinhibition of the oviDN which elicit the oviposition motor action (***Wang et al., 2020***). While this neural pathway is responsible for encoding and conveying mating status (***Yang et al., 2009***; ***Wang et al., 2020***), it does not account for other computations that influence oviposition, such as the evaluations of substrate firmness and nutrient content, which depend on sensory inputs. The neural correlates of these computations are not yet well-defined, but multiple circuits may converge onto the oviposition circuit to influence decision-making and eventually egg-laying behavior.

**Figure 1.**
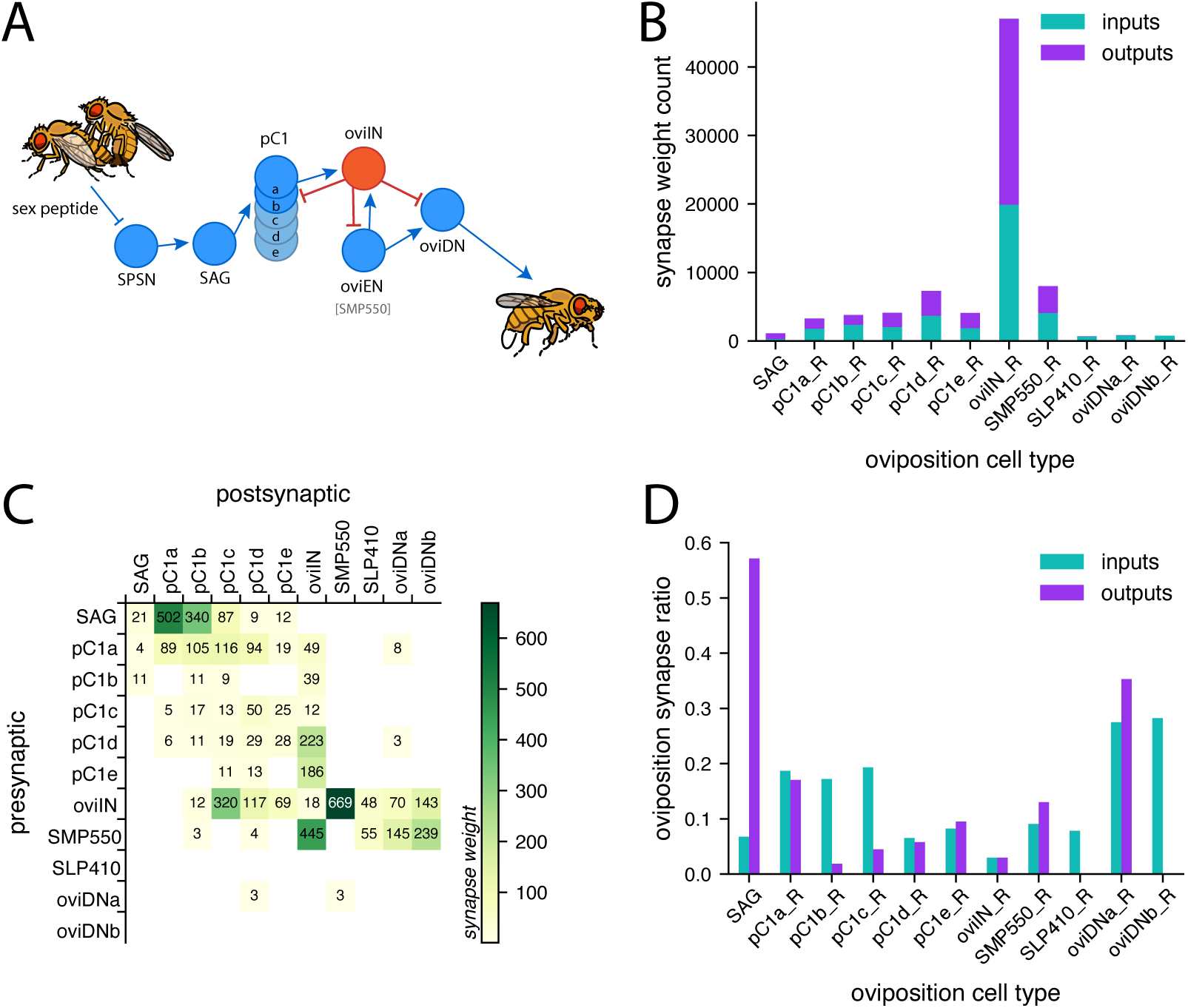
The connectivity profile for the oviposition circuit in the Hemibrain connectome. A. Schematic circuit diagram of the oviposition circuit from ***Wang et al.*** (***2020***) that conveys mating status to the Oviposition Descending Neurons (oviDN) which command the oviposition motor sequence. The oviIN (red) is an inhibitory neuron in this circuit. Fly illustrations are reused and adapted from DataBase Center for Life Science (DBCLS), https://doi.org/10.7875/togopic.2022.325. B. Total synapse counts for the oviposition cell types in the Hemibrain connectome. Counts are based on the number of presynaptic (inputs; teal) and postsynaptic (outputs; purple) synapse connections made by each cell type instance shown (connection weights less than 3 are excluded from the tally). Only right instances of the oviposition cell types are included because left instances are often truncated. The mean counts are presented for the SAG and SLP410_R because they each have two instances. The SMP550 are presumed to be the oviEN, and the SLP410 are believed to be an additional pair of the oviDNa sub-type (***Nojima et al., 2021***). See Figure 1***-1*** for extent of overlap among presynaptic partners. C. Connectivity matrix for the oviposition cell types. Connection weights less than 3 are excluded. Synapse counts are the aggregated synaptic weights from all neurons of one cell type to all neurons of another cell type (combining right and left instances). D. The proportion of synapses that are made with other oviposition neurons. The inputs ratio (teal) for a given cell type is the proportion of synapses received from presynaptic oviposition partners relative to synapses from all presynaptic partners. The outputs ratio (purple) is the proportion of synapses to other oviposition neurons relative to synapses to all postsynaptic partners. Connection weights less than 3 are excluded from this analysis.

In our present study, we begin an exploration of the understudied areas that influence oviposition decision-making by focusing our analyses on the inputs to one central player in the oviposition circuit. Our initial analyses revealed that the oviIN is a highly interconnected neuron in the oviposition circuit and in the brain. We find that the inputs to oviIN predominantly come from the Superior Neuropils of the Protocerebrum (SNP) which include the Superior Lateral Protocerebrum (SLP), the Superior Intermediate Protocerebrum (SIP), and the Superior Medial Protocerebrum (SMP) (***Ito et al., 2014***). These structures have been known to support high-level processing of taste (***Kim et al., 2017***), water-seeking behavior (***Landayan et al., 2021***) that depends on the complex interactions between hunger and thirst (***González Segarra et al., 2023***; ***Jourjine et al., 2016***), locomotor control (***Marquis and Wilson, 2022***), and persistent internal states in female flies (***Deutsch et al., 2020***). Unlike the physical ring attractor within the central complex (CX) or the clearly compartmentalized Mushroom Body (MB), the SNP do not have a structure that reveals their function and are described as a “diffuse” set of neuropils (***Ito et al., 2014***). There is currently no comprehensive map of functional circuits in the SNP, thus, we employed a graph-theoretic approach to identify community structure among the inputs to the oviIN. Our analysis revealed clusters of neurons that preferentially target the oviIN from diffuse regions of the central brain, as well as structured regions. These clusters make anatomically organized synapses on the branches of the oviIN. Ultimately, determining the organization of oviIN’s inputs will facilitate the discovery of novel circuits involved in high-level processing.

## Methods and Materials

### Accessing and working with the Hemibrain connectome data

All analyses presented were performed using the publicly available Hemibrain connectome data released by Janelia (***Scheffer et al., 2020***). The data are from an EM reconstruction of the right hemisphere and partial left hemisphere of a single female fly brain. Morphological data for the 3D reconstruction was accessed, along with tabular data of the neuron annotations and connectivity, using the Neuprint python package (https://github.com/connectome-neuprint/neuprint-python). The data accessed via Neuprint is Hemibrain version 1.2.1. For the brain-wide rankings (***Figure 2***), we relied on the most recent table of traced neurons made available by Janelia (https://storage.cloud.google.com/hemibrain/v1.2/exported-traced-adjacencies-v1.2.tar.gz) which is version 1.2, and which contains only neurons that are traced and uncropped.

**Figure 2.**
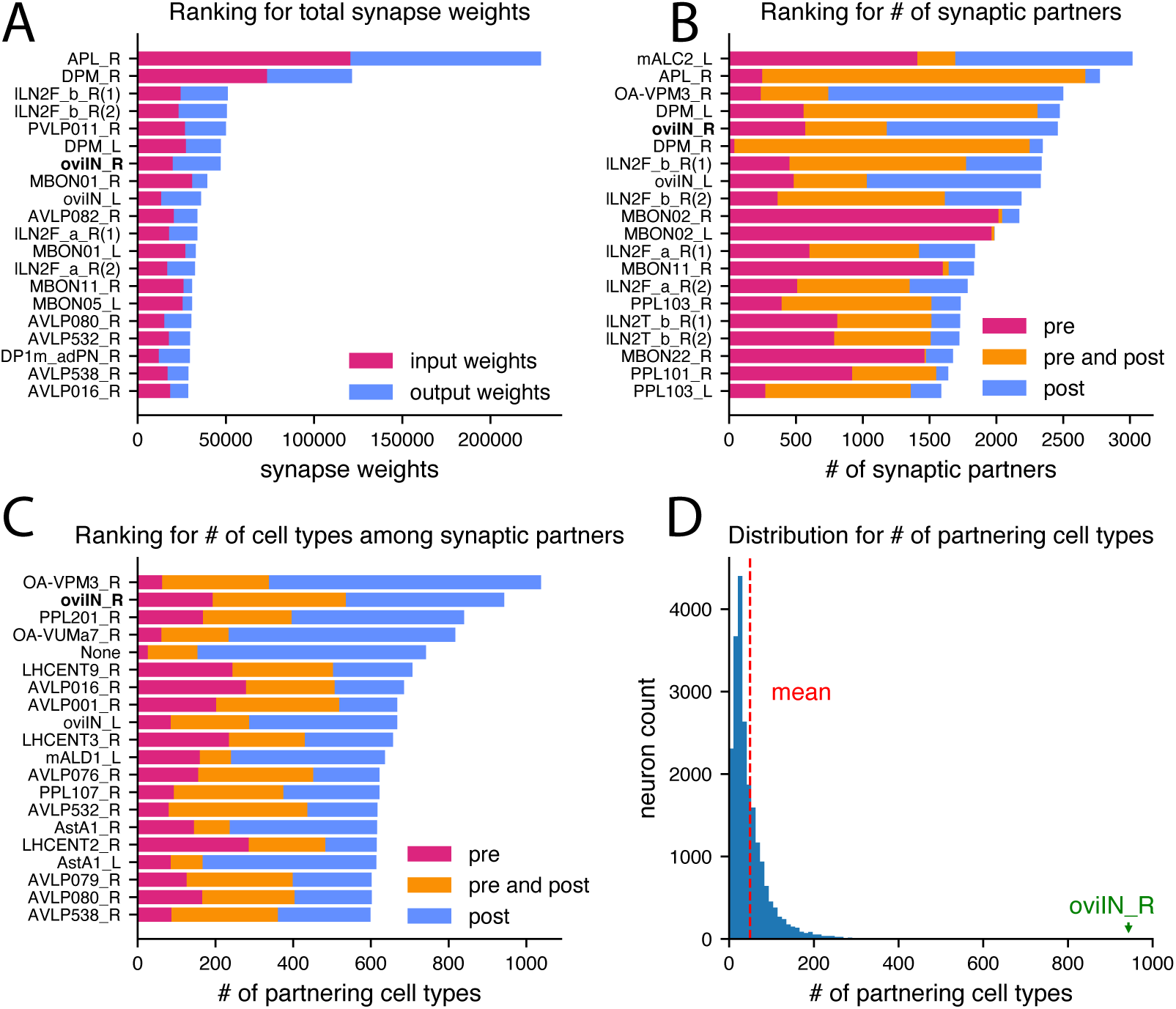
The oviIN is an outlier in the Hemibrain. A. Out of the 21,739 traced, non-cropped neurons in the Hemibrain, the top 20 neurons with the highest total synaptic connections with traced neurons are ranked in descending order and labeled according to their Hemibrain cell type and whether it is a right (R) or left (L) instance (the number in parentheses indicates whether there are multiple instances appearing in the ranking). The right oviIN (oviIN_R) ranks 7th with 47,044 synaptic connections. Input weights (magenta) are counts of postsynaptic sites on the indicated neuron that are paired with presynaptic sites from other neurons. Output weights (blue) are counts of postsynaptic sites that are paired with the presynaptic sites on the indicated neuron. B. The top 20 traced, non-cropped neurons in the Hemibrain with the greatest quantity of unique neuron partners are ranked in descending order and labeled as in (A). Counts of partner neurons that are exclusively presynaptic are shown in magenta, exclusively postsynaptic partners are shown in blue, and counts of neurons that are both presynaptic and postsynaptic to the indicated neuron are shown in orange. Partners making connection weights less than 3 were excluded. The oviIN_R ranks 5th with 2,460 total partners (1,180 presynaptic and 1,890 postsynaptic neurons, of which 610 neurons are both presynaptic and postsynaptic). C. The top 20 traced, non-cropped neurons in the Hemibrain with the greatest quantity of distinct cell types among their direct synaptic partners are ranked in descending order and labeled as in (A). Counts of cell types that are exclusively presynaptic are shown in magenta, exclusively postsynaptic cell types are shown in blue, and counts of cell types that are both presynaptic and postsynaptic to the indicated neuron are shown in orange. Connection weights less than 3 between pairs of neurons were excluded. The oviIN_R ranks 2nd with 943 total cell types among its partners (536 presynaptic and 750 postsynaptic cell types, of which 343 cell types are both presynaptic and postsynaptic). D. Histogram showing the distribution of the number of cell types among synaptic partners for all neurons in the Hemibrain. The average neuron partners with 49 cell types while oviIN_R makes direct connections with 943 cell types.

Every contiguous object in the Hemibrain is assigned a unique bodyId, and many of these are also assigned a cell type. Janelia annotated cell types based on their morphologies and connectivity patterns. There are some traced, non-cropped neurons that are not assigned a cell type. These are labeled as “Nonef” and were excluded from certain analyses wherever indicated. Janelia also annotates neurons with an instance label that can be used to differentiate between right and left instances of bilateral cell types. For example, there are 2 neurons in the Hemibrain of the oviIN cell type: a right and a left instance that are denoted as oviIN_R and oviIN_L, respectively. Our analyses largely focused on the right oviIN.

Neurons in the Hemibrain that match with neurons that have been empirically characterized in the literature are typically annotated with their known name, thus we denote these as characterized cell types. Neurons that have not been characterized are often given a cell type name that begins with the abbreviation for the neuropil where most of its synaptic contacts are located followed by a 3-digit number where the numbering sequence begins with 001. Neurons such as these may have been previously characterized in the literature but have not been matched in the Hemibrain. One such example is SMP550 which is presumed by ***Nojima et al.*** (***2021***) to be the oviEN that ***Wang et al.*** (***2020***) identified in the FAFB volume, but the Hemibrain has not updated its cell type label to reflect that proposition. Except for this particular case, we denote generically-labeled and unnamed cell types (‘None’) as uncharacterized.

*Drosophila* synapses are polyadic, meaning that multiple postsynaptic densities can be associated with a single presynaptic T-bar. The Hemibrain provides counts of the postsynaptic density sites as well as counts of the presynaptic sites on the bodies of anything that is given a bodyId, as well as locations in 3D space of those sites. Additionally, the Hemibrain provides data for the synaptic connections among neurons in the 3D reconstruction. These are given as synaptic weights where the weight is the count of postsynaptic sites within the synapses that connect two neurons. A connection weight between a pair of unique neurons that is less than 3 can be considered a potentially erroneous connection (***Scheffer et al., 2020***). We implement a connection strength threshold of 3 for several of the analyses in our study, wherever indicated.

### Primacy of inputs to oviIN

A neuron or cell type is considered a primary input to oviIN if it has oviIN as its strongest output. Primacy is determined on a by-cell type basis by querying the synaptic outputs of all the neurons of a given cell type, summing those weights by postsynaptic cell type, and putting those aggregated weights in descending order. The position of the oviIN type in the list determines the primacy of oviIN as an output. If oviIN is in the first position of outputs from a neuron, that neuron is a primary input to oviIN (see ***Figure 4***B).

### Parsing neuropil data

When fetching adjacencies between neurons or cell types in the Hemibrain, the data indicates the neuropil, or region of interest (ROI), in which the connection is located. For consistency and clarity, we merged right and left ROIs and relabel primary ROIs with their corresponding supercategory according to ***Ito et al.*** (***2014***) (see ***Figure 5-1***).

For bar chart visualizations in which the data were restricted to synapses made with oviIN (***Figure 4-2***), the synapse weights between oviIN_R and its presynaptic partners were queried by ROI. For each module, the synapse weights from module neurons onto oviIN_R were summed within each neuropil supercategory, thus providing the total synapse weights onto oviIN_R made within each supercategory by the neurons in a given module. For the bar chart visualization in ***Figure 5***A, all presynaptic connections to presynaptic partners of oviIN_R were queried. For each module, the synapse weights from any neuron in the brain onto a presynaptic partner of oviIN_R were summed within each supercategory, thus providing the total synapse weights within each supercategory onto oviIN_R’s presynaptic partners within a given module.

### RenEEL method for computing generalized modularity

We used Reduced Network Extremal Ensemble Learning (RenEEL, ***Guo et al.*** (***2019***)), a machinelearning based approach for network clustering, to identify the partition ∱ which maximizes the generalized modularity density [eqn.1](***Guo et al., 2023***). RenEEL first creates an ensemble of candidate partitions using a fast heuristic. The ensemble is iteratively updated, alternating steps of reducing and replacing. In the reducing step, each community identified by all partitions in the ensemble is combined to a single node. In the replacing step, the fast heuristic is used again on the reduced network. If the result matches one of the previous partitions, or has a lower score than any of the ensemble partitions, the ensemble size is reduced by one. Otherwise, the result replaces the ensemble partition with the lowest modularity score. This process repeats until only one partition remains in the ensemble, representing a consensus best partition. We used an implementation that has previously been shown to produce fast and accurate results for networks of the size and density we consider (***Kunin et al., 2023***; ***Guo et al., 2019***, ***2023***). The source code is available on Github: https://github.com/prameshsingh/generalized-modularity-density.

The set of neurons presynaptic to oviIN_R and the synaptic connections among them were converted into an undirected, weighted graph. The weight of each edge was the total synaptic weight between the two neurons in both directions. The community structure of the network was identified with the partition *C* = {*C*_1_, *C*_2_, …} maximizing the generalized modularity density score (***Guo et al., 2023***),

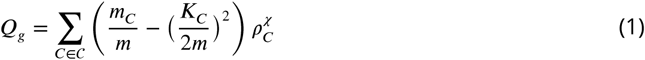

The sum is taken over communities *C*; *m* is the total number of edges in the network, *m*_*C*_ is the total weight of edges in community *C, K*_*C*_ is the weight-degree sum of nodes in *C* (the sum of the weights of edges incident to each node in *C*), and *K*_*C*_ is the relative density of connections in *C*, defined by

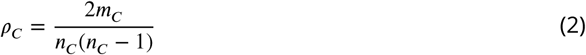

where *n*_*C*_ is the number of nodes in community *C*. The exponent χ controls the resolution of the clustering. At χ = 0, *Q*_*g*_ equals classic modularity density (***Newman, 2006***), with χ *>* 0 resolving smaller, more densely-connected communities (***Guo et al., 2023***).

In our study, RenEEL was applied to the subconnectome of oviIN_R inputs which consists of all the presynaptic partners to oviIN_R and the connections among them. Only traced, non-cropped neurons were included in this subconnectome in keeping with the analyses done by ***Kunin et al.*** (***2023***). To avoid isolating any nodes that could be potentially informative to the clustering, no synaptic weight threshold was applied to the inputs to oviIN_R nor to the connections among those inputs. Thus, the subconnectome was overly inclusive and it contained 1,832 nodes as well as all connections with non-zero weight. The results presented are from a representative run (using χ = 0) that resulted in a single set of partitions, ∱. RenEEL was subsequently run on the same subconnectome 30 times with different random seeds to determine that partitions were largely consistent between runs (mean Jaccard similarity: 0.89). The weak inputs to oviIN drove most of the variability in the results while the primary strong inputs to oviIN were more consistently partitioned. When analyzing and making comparisons to the modularity of the full Hemibrain, RenEEL results from ***Kunin et al.*** (***2023***) were used.

### Analyses and Visualizations

Code Accessibility

All analyses and visualizations were done in Python using custom-written scripts along with various published and open-source packages. Our analysis code is publicly available on GitHub: https://github.com/Gutierrez-lab/oviIN-inputs. We used Mac OS computers for most work, and a High Performance Computing Cluster for running the RenEEL algorithm on the connectome data.

### Joint clustering plots

We compared the modularity results for the oviIN_R inputs subconnectome clustering and the full Hemibrain connectome clustering. We computed the proportion of nodes from a module in one clustering that are partitioned into a module from the other clustering. The width and height of the boxes in ***Figure 5***B represent the joint counts of neurons within two modules over the total counts within a module of the respective dataset. If N*_Hi_* is the number of nodes in module *i* of the full Hemibrain clustering, and N*_O_*_j_ is the number of nodes in module *j* of the oviIN_R input clustering, and the number of nodes jointly appearing in module *j* of the full Hemibrain clustering and in module *i* of the oviIN_R input clustering is 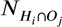, then the width of any box is 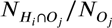 and the height of any box is 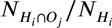.

### Jaccard Similarity

To evaluate the similarity between different neuron clusterings *A* and *B*, we compute the Jaccard similarity. For a set of neurons {1, 2, 3, … *N*}, Jaccard Similarity considers all pairs of neurons in the set and is defined as:

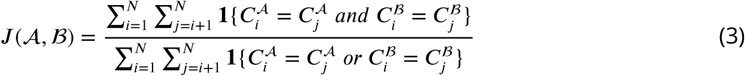

where *N* is the total number of neurons in the set, 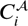 and 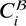 are the cluster assignments of neuron *i* in clusterings *A* and *B*, respectively. The indicator function **1**{⋛} equals 1 if the condition inside is true and 0 otherwise.

The Jaccard similarity ranges from 0 to 1. A value of 0 indicates no overlap between the clusterings, while a value of 1 indicates perfect overlap. Intermediate values indicate that some pairs of neurons are clustered together for both clusterings. In our study, we applied this method to compare the clustering of the oviIN_R inputs with the clustering of the entire Hemibrain for different resolutions of RenEEL algorithm.

### Input Similarity

Input similarity was computed as the cosine distance between the vectors representing the inputs to each neuron. Subconnectome neurons were removed from the inputs to the subconnectome neurons. For every neuron *n*, we compute the vector *n_in_* of length equal to the number of neurons in the Hemibrain, with the *i*th entry equal to the synapse weight from neuron *i* to neuron *n*. If neuron *i* is not synaptically connected to neuron *n*, the corresponding entry is 0. For two vectors *u, v* the cosine distance is given by 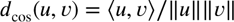. Because all entries in *n_in_* are nonnegative, cosine similarity ranges between 0 and 1. A similarity of 0 means that two neurons receive inputs from completely disjoint sets of neurons; a similarity of 1 means they receive inputs from exactly the same neurons in exactly the same proportion.

Mean similarity was computed within modules by averaging over all pairs of distinct neurons belonging to that module, and across modules by averaging over all pairs with one neuron in each cluster. For comparison, we ran 100 trials with shuned module membership, preserving the size of each module but randomly assigning module labels to each neuron. The z-scores were computed from the distributions of mean pooled similarity of the shuned data relative to the mean pooled similarity of the true data.

## Results

### oviIN is an outlier in the oviposition circuit

We queried the Hemibrain (***Scheffer et al., 2020***) for the oviposition cell types from ***Wang et al.*** (***2020***) (***Figure 1***A) and found the quantity of presynaptic and postsynaptic connections made by each of the oviposition cell types (***Figure 1***B). The left instances are sometimes truncated and less complete than the right instances, thus left instances of oviposition cell types were excluded from the results. We were immediately struck by the oviIN because it makes far more synaptic connections than the other oviposition cell types (***Figure 1***B). Particularly relevant is the quantity of input connections to oviIN. Even when excluding connection strengths less than 3, the right-hemisphere oviIN (oviIN_R) has 19,882 synaptic input connections which is far more than the quantity of inputs received by any other oviposition neuron.

The oviIN makes abundant connections within the oviposition circuit (***Figure 1***C), particularly with SMP550 which is presumed to be the oviEN (see ***Nojima et al.*** (***2021***)). The oviIN and oviEN are strongly, reciprocally connected. As expected, oviIN is presynaptic to all subtypes of oviDN (including the putative additional pair of oviDNa that are labeled as SLP410 in the Hemibrain; see ***Nojima et al.*** (***2021***)) where it inhibits the initiation of the oviposition sequence. Among the pC1 neurons, oviIN receives inputs from all subtypes a-e and also sends outputs to subtypes b-e, forming a small recurrent motif. In accordance with whole-cell recordings from ***Wang et al.*** (***2020***), the SAG neurons strongly contact the pC1a-b neurons. SAG neurons also make moderately strong connections to pC1c in the Hemibrain data. The pC1a-b neurons make moderate connections onto the oviIN. Interestingly, the oviIN provides some feedback connections to the pC1b neurons (the 1 synaptic connection to one of the pC1a neurons was below the synaptic strength threshold of 3 for ***Figure 1***C). The remaining pC1 neurons are not among the subtypes that are believed to directly convey mating status to the oviposition neurons, but as noted elsewhere, the pC1 neurons are recurrently connected and may share mating status information received by the pC1a-b neurons (***Wang et al., 2020***; ***Deutsch et al., 2020***). We note that pC1c sends relatively weak connections to oviIN but receives relatively strong feedback from oviIN. The connections that pC1d-e neurons make onto oviIN are stronger than those from pC1a-b onto oviIN. Although the pC1d-e neurons receive input from SAG neurons, they were not found to be responsive to SAG activation by ***Wang et al.*** (***2020***). The pC1d-e neurons also receive feedback from oviIN.

While oviIN forms strong synaptic connections with many of the oviposition cell types (we define strong synaptic connections among cell types as ≤ 100 connection weights), overall connections with the oviposition neurons make up a very small proportion of oviIN’s synapses (***Figure 1***D). Roughly 3% of synapses on oviIN_R are from other oviposition neurons. It is clear from the Hemibrain data that oviIN makes a vast number of connections outside of the oviposition circuit. Furthermore, there is little overlap among the inputs that oviIN_R receives and those received by the right instances of oviEN (SMP550) and the oviDN subtypes (***Figure 1-1***). If we only consider connections with a weight of 3 or greater, we find that oviIN_R and SMP550_R have 37 presynaptic partners in common, which is a small portion of the presynaptic partners to SMP550_R (235) and to oviIN_R (1180). Only 3 of those 37 overlapping partners are cell types making strong inputs to oviIN_R. Since oviIN makes so many of its connections outside of the oviposition circuit, and it receives inputs that are largely distinct from those received by oviEN and oviDN, we chose to focus on analyzing the connectivity of oviIN – particularly its inputs.

### oviIN is an outlier in the brain

Given that oviIN’s extensive connectivity makes it an outlier in the oviposition circuit, we next sought to determine whether it is an outlier among the rest of the neurons in the Hemibrain with an eye towards other large inhibitory neurons. To eliminate possibly erroneous connections, we excluded connection weights of less than 3 between pairs of neurons from consideration in the subsequent rankings. We found that the oviIN is one of the most synaptically endowed neurons in the Hemibrain. The right oviIN is the 7th highest ranking neuron for total synaptic connections (47,044 synapse weights; ***Figure 2***A, ***Figure 1***B). The Anterior Paired Lateral (APL) neuron has the most synaptic connections (228,713 synapse weights) out of all the 21,739 traced, uncropped neurons in the Hemibrain dataset. The APL has a role in normalizing the activity of odor-encoding Kenyon cells in the MB (***Prisco et al., 2021***; ***Pitman et al., 2011***). In the synaptic connection ranking, it is closely followed by the right Dorsal Paired Medial (DPM) neuron which is involved in odor memory and associative learning (***Waddell et al., 2000***; ***Keene et al., 2004***, ***2006***), and is sleep-promoting (***Haynes et al., 2015***). The local Lateral Neurons (lLNs) of the antennal lobes also rank highly for synaptic connections. Notably, these highly synaptically-endowed neurons are large inhibitory neurons (***Haynes et al., 2015***; ***Pitman et al., 2011***; ***Chou et al., 2010***). The average neuron in the Hemibrain makes a total of 1,198 synaptic connections. When considering the extent of oviIN_R’s inputs separately, it ranks 14th for synaptic input connections (19,882 synapse weights), making it an outlier in terms of the amount of input it receives.

The right oviIN ranks 5th for the number of total synaptic partners (pre and post) with 2,460 distinct partnering neurons (***Figure 2***B), which is well above the mean number of partners for neurons in the Hemibrain (111). APL ranks 2nd with 2,775 partners, while the DPMs and lLNs have 2,411 and 2,264 on average, respectively. When considering only presynaptic partners, oviIN ranks 23rd while APL ranks 1st. However, the oviIN_R receives inputs from 1,180 neurons in the Hemibrain while the average neuron in the Hemibrain only receives inputs from 66 neurons.

The oviIN distinguishes itself from these other large inhibitory neurons with the number of cell types that it interacts with. With 943 cell types among its synaptic partners, oviIN ranks 2nd in the Hemibrain on that metric (***Figure 2***C). Despite a high ranking for the total number of synaptic connections, APL only ranks 235th in terms of the number of cell types among its direct connections with 250 cell types. The DPMs and lLNs connect with 90 and 219 cell types on average, respectively, while the average neuron in the Hemibrain makes direct connections with 49 cell types (***Figure 2***D). Notably, the oviIN_R ranks 1st when only taking into account the number of cell types that are presynaptic to it. The oviIN_R receives inputs from 536 cell types while neurons in the Hemibrain on average receive inputs from 27 cell types. Compared to the average neuron in the Hemibrain which makes connections in 2.6 of the 12 neuropil supercategories, oviIN_R connects twice as many. The right oviIN makes synaptic connections in 5 supercategories (***Figure 5-1***) with most of those connections occuring in the SMP.

### “Hub-and-bespoke” structure among inputs to oviIN

One could imagine that oviIN’s extensive connectivity might be a result of it receiving small amounts of input from many different neurons. On the contrary, we find that oviIN’s strongest presynaptic partners also favor oviIN as a postsynaptic partner (***Figure 3***A). The oviIN_R is the primary target for the majority of cell types that strongly synapse onto it (with ≤100 synapse weights; 56.25%). As might be expected, this primacy decreases for cell types that do not send strong connections to oviIN, indicating that those cell types prioritize making connections with other neurons besides oviIN. As a point of comparison, a much smaller portion of the strong inputs to the APL primarily send their synapses to APL (4.3%). Only the LHMB1 and LHCENT5 cell types are primary strong inputs to APL. This finding confers with APL’s divisive normalization function where it samples from many neurons to obtain a population-level average of activity. As such, the oviIN’s role does not appear to be divisive normalization. Rather, the primacy of the strong inputs to oviIN is suggestive of dedicated pathways of information flow.

**Figure 3.**
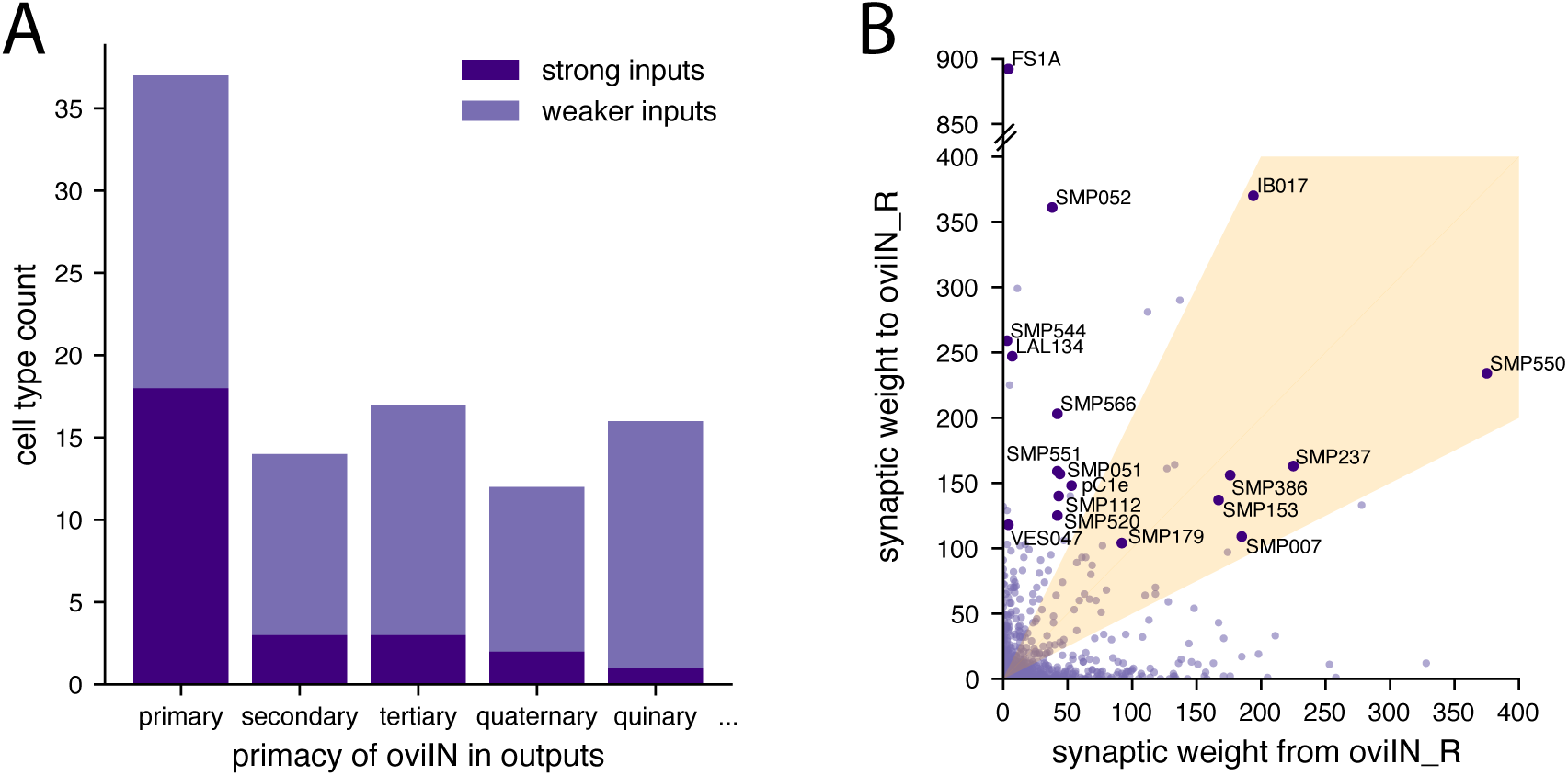
A. Cell types that have oviIN as their primary output also tend to be strong inputs to oviIN. Strong inputs make ≤100 synaptic connections to oviIN_R (dark purple); weaker inputs make *<*100 synaptic connections to oviIN_R (light purple). There are 37 cell types that have oviIN as their primary output. Among those, 18 cell types make 100 or more synaptic connections to oviIN_R. Cell types making strong connections to oviIN_R make up a smaller proportion of cell types that most strongly output to another cell type besides oviIN. B. Each point on the graph represents a cell type that is a direct synaptic partner of oviIN_R. The combined synaptic weight from oviIN_R to a cell type is plotted along the x-axis while the combined synaptic weight to oviIN_R from a cell type is plotted along the y-axis. The 18 primary strong inputs to oviIN_R (dark purple points) are annotated with their cell type label. A subset of the primary strong inputs to oviIN_R occupy the zone of high reciprocity with oviIN_R (orange cone) where connections in the stronger direction are no more than twice the strength of connections in the weaker direction.

The primary strong inputs to oviIN lay the foundations for potential pathways to oviIN from other circuits, and their identities may be crucial for determining which circuits they funnel information from. However, the majority of cell types connecting to oviIN are uncharacterized, meaning they have either not been studied before being identified in the Hemibrain or they have not yet been identified from existing scientific literature. The Hemibrain provides generic labels to uncharacterized cell types that indicate the neuropil in which the bulk of their synapses are found (***Scheffer et al., 2020***). Neurons that have similar morphology and significant overlap in their synaptic connection patterns are assigned to the same cell type in the Hemibrain. A large proportion of the inputs to oviIN, including its primary strong inputs, are cell types with Hemibrain-provided generic cell type labels that indicate that they mainly reside in the Superior Medial Protocerebrum (SMP). However, oviIN’s strongest input comes from the FS1A cell type. The FS1A neurons are a columnar output cell type of the Fan-shaped Body (FB) (***Hulse et al., 2021***). The FS1A neurons mainly receive FB inputs from layers 2-6 and leave the FB to project outputs to the SMP and Crepine. The pC1d and pC1e are also among the strongest inputs to oviIN. The pC1d has been shown to promote aggressive behavior in the female fly (***Schretter et al., 2020***), while activation of pC1d-e together drives a persistent response of several minutes indicative of an internal state encoding (***Deutsch et al., 2020***).

Among the primary strong inputs to oviIN, some of the cell types form highly reciprocal connections with oviIN_R while others have more of a feedforward relationship with it (***Figure 3***B). The FS1A, SMP544, LAL134, and VES047 cell types hardly make any feedback connections to oviIN_R. In contrast, the SMP179, SMP386, SMP153, SMP237, SMP550, SMP007, and IB017 cell types are strongly reciprocal with oviIN_R. These findings imply that FS1A, SMP544, LAL134, and VES047 neurons directly convey information from other circuits while the information conveyed by oviIN’s reciprocally connected primary strong inputs, which are primarily uncharacterized SMP types, can be modulated by oviIN.

Clustering the inputs to oviIN

The FS1A and pC1 neurons are not known to have any overlapping functions, suggesting that oviIN integrates information from at least two distinct circuits. However, the right oviIN receives inputs from 943 cell types leading us to think that many of those cell types might be associated with the same circuits. Furthermore, there is the problem of what to make of the hundreds of uncharacterized cell types among the inputs to oviIN. To begin to organize the inputs projecting to oviIN, we used a network community detection method. We employed a machine learning algorithm, RenEEL (***Guo et al.*** (***2019***); see Methods and Materials), to find clusters of densely connected neurons from an undirected network where the nodes are neurons and the links are weighted synaptic connections.

RenEEL maximizes modularity which is a measure of the excess of within-group connections compared to a randomly-connected network (***Newman, 2006***). The goal of maximizing modularity is to find an optimal partitioning of the network into modules (i.e. clusters) with higher within-group than between-group connectivity. Generalized Modularity analysis is a multi-resolution method that can find community structure at various scales by adjusting a resolution scale parameter (***Guo et al., 2023***). At a low resolution, the coarse structure of the network is returned as a small number of modules with many neurons in each. As the resolution is increased, the network is partitioned into a greater number of smaller modules. The clusters of densely connected neurons identified in this way may form the basis of a hierarchical circuit structure. Previous studies identified community structure in the full Hemibrain connectome at multiple resolutions by maximizing Generalized Modularity Density [eqn.1] (***Kunin et al., 2023***). This unsupervised analysis recovered the gross structure of the brain by returning modules that reflect the neuropil super-structures. At higher resolutions, known fine structures, such as the sub-circuits of the layered FB, were recovered.

In the current study, we limit our analysis to classic modularity (coarse resolution) of the network of inputs to the right oviIN. Since this network is a part of the full connectome, we refer to it as a subconnectome. The subconnectome of inputs to oviIN_R contains all of oviIN_R’s presynaptic partners and the connections among them, and it excludes all other connectome neurons. Our goal was to identify community structure that may otherwise be subsumed by the superstructure of the neuropils when analyzing the full Hemibrain connectome. The modular structure we uncover depends on the relative connection strengths within and between groups; thus, by eliminating neurons that do not form direct connections to oviIN_R we can hope to identify denselyconnected communities that potentially span multiple brain areas. RenEEL returned 7 modules containing from 199 to 347 neurons each for the subconnectome of inputs to oviIN_R (***Table 4-1***. Each module contains between 111 and 199 unique cell types.

### The modules divide the primary strong inputs to oviIN_R

The most prominent inputs to oviIN_R are spread out among the different modules (***Figure 4***A), indicating that they represent parallel pathways that funnel distinct information to oviIN_R. The FS1A cell type is clustered into the same module as the other FB cell types that send strong inputs to oviIN (FC2B and FC2C; module 4) along with select, uncharacterized SMP cell types. The pC1d-e are clustered together in module 3. The primary strong inputs to oviIN_R are also spread among the modules (see primary bar in ***Figure 4***B), supporting the notion that if oviIN is a hub, the primary strong inputs define the spokes that convey information from distinct circuits.

**Figure 4.**
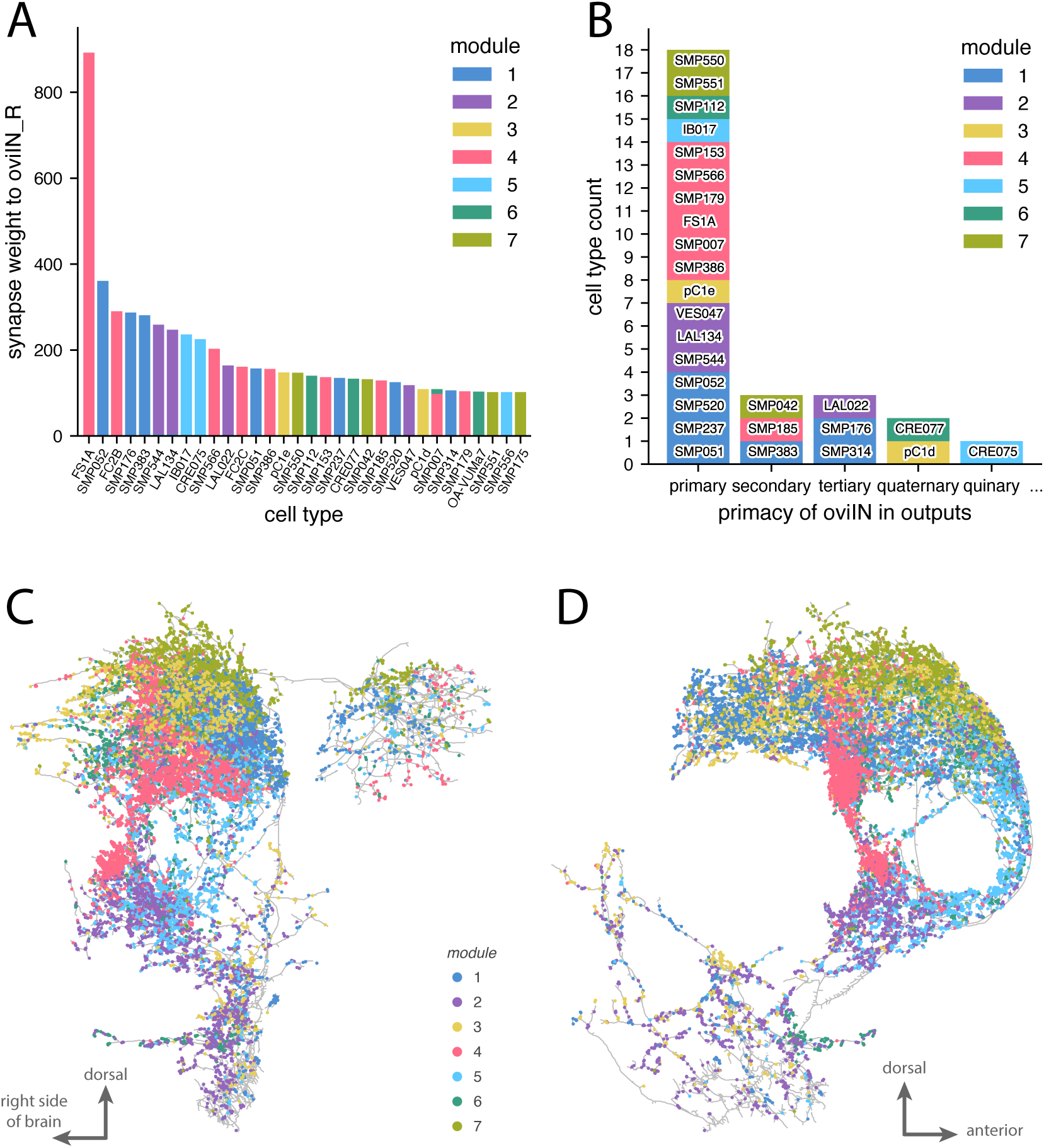
Modularity maximization reveals mesoscale and microscale connectome structure among inputs to oviIN_R. A. The strongest inputs to oviIN_R. Neuronal cell types that make an aggregated synapse count of at least 100 onto oviIN_R are shown along the x-axis. The bars indicating synapse counts are color-coded by the module(s) into which the neurons of a type are clustered. Sometimes, neurons of the same type are clustered into different modules. For example, three instances of SMP007 are grouped into module 4 and the other instance is grouped into module 6 (there are four instances of SMP007, all on the right side of the brain). B. Most of the strongest inputs to oviIN_R are primary inputs, meaning that the cell type primarily targets oviIN. Primacy indicates oviIN_R’s ranking among the cell type’s strongest outputs. Only cell types with aggregated connection strength ≤ 100 onto the oviIN_R are shown. Each cell type is colored according to the module where most of its instances are clustered. C. Front view of synaptic input sites on oviIN_R colored by module. The reconstructed skeleton of the right oviIN from the Hemibrain is outlined in gray. Each dot on the body of oviIN_R is a postsynaptic site. These connection sites are colored according to the module of the presynaptic neuron. Synapses from neurons without a cell type label (annotated as ‘None’) are not shown, but results are qualitatively the same when they are included. See Figure 4***-1*** for individual modules, ***Table 4-1*** for module membership, and Figure 4***-2*** for neuropil supercatergories where synapses are located. D. Lateral view (from the right side of the brain) of synaptic input sites on oviIN_R colored by module as in (C).

Although most of the cell types in these modules are uncharacterized, there are a few characterized cell types that corroborate the potential functional basis of the modules. The aIPg1-3 cell types were previously found to elicit aggressive behavior from female flies and to form a recurrent network with the pC1d-e (***Schretter et al., 2020***; ***Deutsch et al., 2020***). The aIPg cell types make moderate connections to oviIN, and they are clustered together with the pC1d-e in module 3 (***Table 4-1***). We note that pC1a-b are clustered into module 7 with the oviEN (SMP550) while the pC1c are in module 1. Module 7 has other *fru+* (fruitless-expressing) neurons that, along with the pC1ab neurons, are broadly implicated in female receptivity. These include vpoDN (***Wang et al., 2021***), SAG (***Feng et al., 2014***), and a non-fruitless cell type, SMP286 (a.k.a. pCd-2; ***Imoto et al.*** (***2024***)). Also in module 7 is the SMP029 which is a candidate for the aDN (***Matsliah et al., 2023***) which have a role in site selection during oviposition (***Nojima et al., 2021***).

These examples are only a handful of neurons and cell types from among the hundreds of neurons in their respective modules and therefore it would not be reasonable to ascribe a specific function to the entire module based on the minority of cell types that happen to be characterized. However, it is reassuring that characterized neurons with related functions were found in the same module. Furthermore, the relationships between characterized neurons and the primary strong inputs to oviIN in their module may provide clues about the functions of those uncharacterized primary strong inputs.

The only clock neurons that directly connect to oviIN are two LPN and two instances of the LNd cell type which correspond to the E1 subtype of evening cells (***Shafer et al., 2022***). These neurons are clustered together in module 1 with aMe24, a cell type from the accessory medulla, an area involved in circadian rhythms (***Helfrich-Förster et al., 2007***). Module 1 is also where the left oviIN was found. The E1 LNds have reciprocal connections with the SMP520 which is a primary strong input to oviIN that was clustered into module 1 (***Figure 4***B). Whether or not the SMP520 carries information about circadian timing to oviIN remains to be seen because the LNd inputs (indeed, any clock inputs) to SMP520 are not among its strongest inputs. However, three other primary strong inputs from module 1, SMP237, SMP051, and SMP052, make a small, strongly recurrent network with the aMe24 and pC1c.

Modules 5 and 6 are peppered with mushroom body output neurons (MBONs). The three MBON cell types in module 5 (MBON27, MBON33, MBON35) are atypical MBONs, meaning they receive inputs from outside the MB (***Li et al., 2020***). MBON27 is specialized for visual inputs (***Li et al., 2020***), and it is implicated in a small network connecting CX to MB along with oviIN and mALD1 which appeared in module 5 as well (***Hulse et al., 2021***). The sole primary strong input to oviIN_R from module 5, IB017, makes relatively weak reciprocal connections with mALD1 and barely connects to the MBON27, indicating that it is not part of that CX to MB circuit. However, IB017 does form moderate connections with MBON35 and MBON33. The MBONs in module 6 are a mixture of typical and atypical types (MBON01, MBON04, MBON05, MBON12; atypical: MBON26, MBON31, MBON32). The connectivity profile of the MBON12 suggests that it has a role in water-seeking behavior, and MBON26 receives a significant amount of thermo-hygrosensory input (***Li et al., 2020***). The SMP112, the only primary strong input to oviIN_R in module 6, does not receive strong inputs from MBON12 and MBON26 but it does receive moderate input from MBON31 and MBON32 which are the strongest inputs to MBON26.

As mentioned, the FB cell types that project to oviIN_R were mainly clustered into module 4. The FS1A, FC2B, and FC2C form a strongly recurrent network. The SMP cell types in module 4 that are primary strong inputs to oviIN_R make very sparse connections to these FB cell types, but the SMP386, SMP153, and SMP007 receive moderate to strong inputs from the FB cell types, indicating that the flow of information goes from the FB neurons to the recurrently connected network of oviIN, SMP386, SMP153, and SMP007. In particular, oviIN and FS1A are the strongest cell type inputs to SMP386. FS1A is the strongest primary input to oviIN, but the SMP386, SMP153, and SMP007 are also primary strong inputs to oviIN and they may form a secondary pathway through which FB information can reach the oviIN, or they may form a circuit that performs a distinct computation on FB inputs before conveying the results to the oviIN.

### Anatomical organization of the modules

RenEEL was run 30 times on the same subconnectome data with different random seeds and it largely returned the same modules. While this gives us confidence in the RenEEL algorithm’s ability to detect fairly consistent communities within the network of oviIN_R’s inputs, it does not on its own confirm that RenEEL has uncovered biologically-relevant brain structure. To determine whether there is any spatial organization inherent in the modules, we visualized the input synapses to oviIN_R on its reconstructed skeleton (***Figure 4***C,D; ***Figure 4-1***). The input synapses on oviIN_R are shown as dots and each dot is colored according to the module of the presynaptic neuron. Neurons from modules 1, 3 and 7 mostly synapse onto oviIN_R in the dorsal tuft of processes within the SMP where most of oviIN_R’s output sites also reside (***Figure 4-1a***, ***Figure 4-1c***, ***Figure 4-1g***). Neurons from modules 4 and 6 also form most of their synapses onto oviIN_R in the dorsal tuft, but they also have a large portion of synapses in the inferior neuropils along the posterior and anterior part of the Crepine, respectively (***Figure 4-1d***, ***Figure 4-1f***). Module 5 neurons predominantly form their synapses to oviIN_R along the anterior Crepine, but also occupy portions of the dorsal tuft (***Figure 4-1e***). Neurons from module 2 mainly form their connections along the ventral portion of the oviIN_R intersecting with lateral complex (LX), ventromedial (VMNP) and inferior neuropils (INP) (***Figure 4-1b***). The modules do not simply reflect the boundaries of the neuropils that intersect with oviIN_R because all of the module synapses intersect with a mixture of neuropils and they all intersect to some extent with the SNP (***Figure 4-2***). Nonetheless, the anatomical organization of modules indicates that RenEEL found biophysically meaningful clusters of neurons. The organization along the distinct branches of oviIN_R suggests that neurons work together with the other neurons in their module to influence the activity within different regions of this large neuron. It is possible that the motifs within these modules act as local processing units to drive, summate, or shunt activity from other local processing units.

It is reasonable to wonder whether the anatomical organization of the modules is a reflection of the neuropil organization of the brain more broadly, rather than the neuropil boundaries of the synapses on oviIN_R. In other words, has RenEEL clustered oviIN_R’s inputs according to the neuropils that they fibelong tofi? Strictly speaking, neurons cannot easily be classified as belonging exclusively to one neuropil because they often make connections in multiple neuropils, although researchers sometimes classify a neuron based on the neuropil where it makes the majority of its synaptic contacts (***Scheffer et al., 2020***; ***Schlegel et al., 2024***; ***Ito et al., 2014***). To determine the various brain regions that are represented among the inputs to oviIN_R, we queried the Hemibrain for the neuropils in which oviIN_R’s presynaptic partners receive synaptic connections from any neuron. We found that RenEEL did not simply partition oviIN_R’s input connectome according to the neuropils where those neurons receive their connections (***Figure 5***A). In all but modules 2 and 5, the neurons receive the majority of their synapses in the SNP. We had already determined that, collectively, the majority of input connections to oviIN_R are made in the SNP (see ***Figure 4-2***), and that a large proportion of oviIN_R’s input connectome contains neurons that are labeled as SMP neurons. ***Figure 5***A shows that RenEEL has identified SNP sub-groups that may pertain to distinct circuits in the SNP. Neuropil designations are primarily driven by anatomical considerations and do not necessarily demarcate functional circuits (***Ito et al., 2014***; ***Yu et al., 2013***). Thus, the modules may loosely represent circuit structure that draws from SNP together with other neuropils. For example, the most prominent cell type input to oviIN, FS1A, is in module 4. The FS1A neurons receive most of their input connections in the FB neuropil of the CX, but they make most of their output connections in the SMP (of the SNP) and Crepine (of the INP) (***Hulse et al., 2021***). Cell types such as this may form a bridge from one part of the brain to the SNP, INP, LX, or VMNP neuropil supercategories where they interact with oviIN.

**Figure 5.**
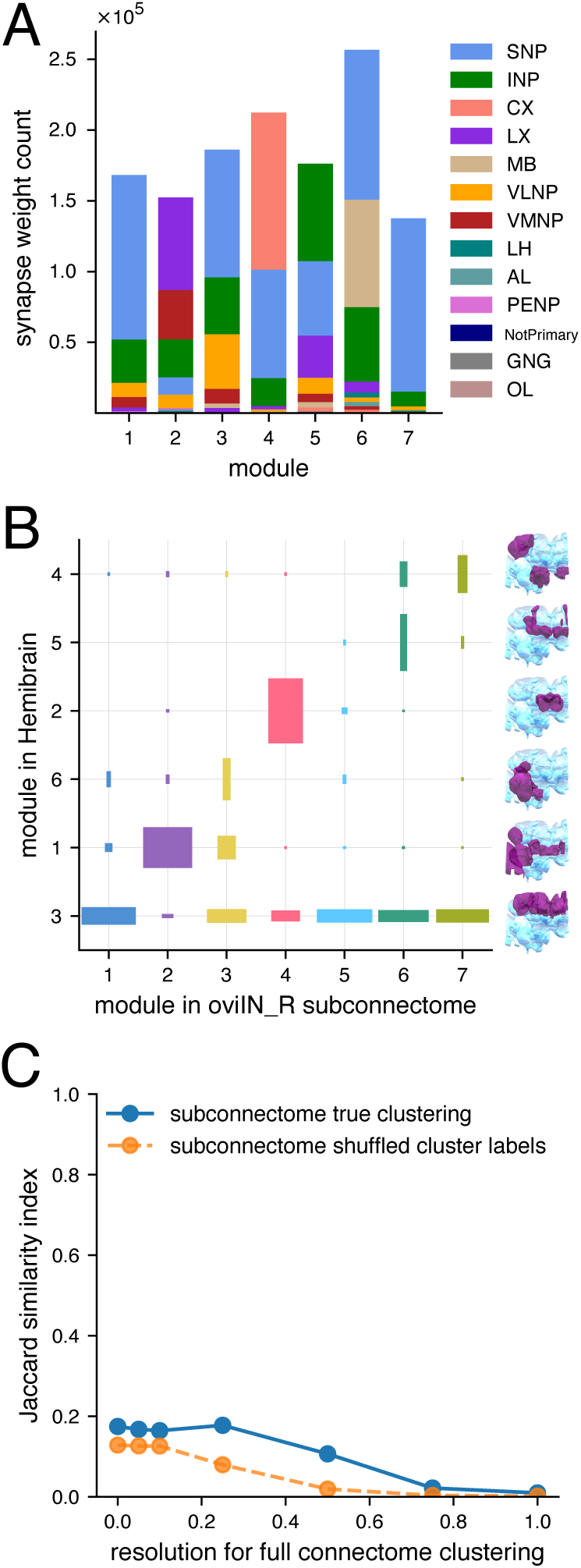
A. The counts of connections received within the various neuropil supercategories by the neurons collectively in each module. The neurons in most of the modules receive a large portion of their synaptic input in the SNP. We use the supercategories defined by ***Ito et al.*** (***2014***) (see Figure 5***-1***). B. Modularity maximization returns partitions of the oviIN_R input subconnectome that are distinct from partitions for the full Hemibrain connectome. The seven modules that we found for the subconnectome of oviIN_R’s inputs are shown along the x-axis. The modules for the full Hemibrain found by ***Kunin et al.*** (***2023***) are on the *y*-axis, and an anatomical rendering is provided for each of those modules on the righthand side of the plot showing their most closely corresponding neuropils (purple) within the brain (light blue). The width and height of each box is an indication of the portion of nodes that jointly appear in a given module of the subconnectome clustering and a given module of the full connectome clustering. The colors correspond to the subconnectome clustering from Figure 4. C. Jaccard similarity between the coarse clustering of the subconnectome of oviIN_R’s inputs and the full connectome at various clustering resolutions (solid blue line), from coarse (0.0 resolution) to fine (1.0 resolution). Jaccard similarity for a random baseline is plotted in orange (dashed line) where the cluster labels for the subconnectome were randomly shuffled before being compared to the full connectome clustering.

### Functional organization of the modules

We next wondered whether the partitions found for oviIN_R’s input connectome reflect those previously found for the full Hemibrain connectome by ***Kunin et al.*** (***2023***). The coarse modules they found in the full Hemibrain connectome were roughly aligned with the neuropil structure of the brain, indicating that RenEEL identified macroscale brain structure. Even though we only consider neurons which make direct synaptic contact with the right oviIN in the present study, we might expect strong alignment with the full Hemibrain modules if the brain’s super-structure is still present within the subconnectome of oviIN_R’s inputs. We compared our results to the modules identified by ***Kunin et al.*** (***2023***), and we did not find strong alignment. The portion of oviIN_R’s input connectome that resides in full Hemibrain module 3 – which corresponds mostly to SMP and SIP neuropils – is spread out among all of the subconnectome modules (***Figure 5***B). In other words, large portions of the neurons in the subconnectome modules all share a common module in the full Hemibrain clustering – namely the module that encompasses the medial and intermediate superior neuropils. Full Hemibrain module 1 corresponds mainly to the VMNP, and the portion of oviIN_R’s connectome that is from that area is dominated by subconnectome module 2. Likewise, full Hemibrain module 2 (which corresponds to FB) is mainly represented in subconnectome module 4. However, all of the subconnectome modules are composed of a combination of the full Hemibrain modules. Since every cluster identified in oviIN_R’s connectome has significant overlap with the full Hemibrain module that encompasses the SIP and SMP, this suggests that, in the full Hemibrain, the connections within SIP and SMP are stronger overall than connections between these areas and other neuropils, and that connections within potential circuits that include SIP and SMP neurons combined with other cell types are dense enough to be detected in the absence of the connections with the broader network of each neuropil.

To determine whether the coarse modules of oviIN_R’s input connectome correspond to mid-resolution modules of the full Hemibrain, we computed the Jaccard similarity [eq.3] between the coarse clustering of the subconnectome of oviIN_R’s inputs and the clusterings of those same neurons in the full connectome at various resolutions. A high Jaccard similarity (close to 1) is an indication that many pairs of neurons are partitioned into a module together in both the subconnectome clustering and the full connectome clustering at a particular resolution. We found low similarity between the subconnectome clustering and the full connectome clustering (***Figure 5***C). For all resolutions of the full connectome clustering, Jaccard similarity with the subconnectome clustering was less than 0.2. However, Jaccard similarity was always slightly higher than a random baseline in which the module labels for the subconnectome were shuned. Thus, applying RenEEL to a subconnectome of the brain, as we have done for oviIN_R’s input connectome, does not partition the neurons in the same way as for the full connectome, either at the coarse or higher resolutions. Our results suggest that the modules correspond to a microscale organization at the level of oviIN_R’s inputs – which may correspond to distinct circuit pathways and which may have functional significance for oviIN.

To verify that these modules are representative of distinct pathways that converge onto oviIN_R, we once again considered their connections within the full Hemibrain. We computed the similarity of inputs from anywhere in the brain to each pair of neurons in oviIN_R’s input connectome (see Methods and Materials). On average, the similarity of within-module pairs exceeds those of between-module pairs (***Figure 6***A). In other words, presynaptic partners of oviIN_R have more of their inputs in common with other neurons in their same module than with neurons in a different module. Moreover, comparison to a randomly partitioned network reveals that within-module pairs are more similar than would be seen by chance (***Figure 6***B, ***Figure 6-1***). Taken together, these results support the idea that the modules approximate parallel streams of input from different circuits.

**Figure 6.**
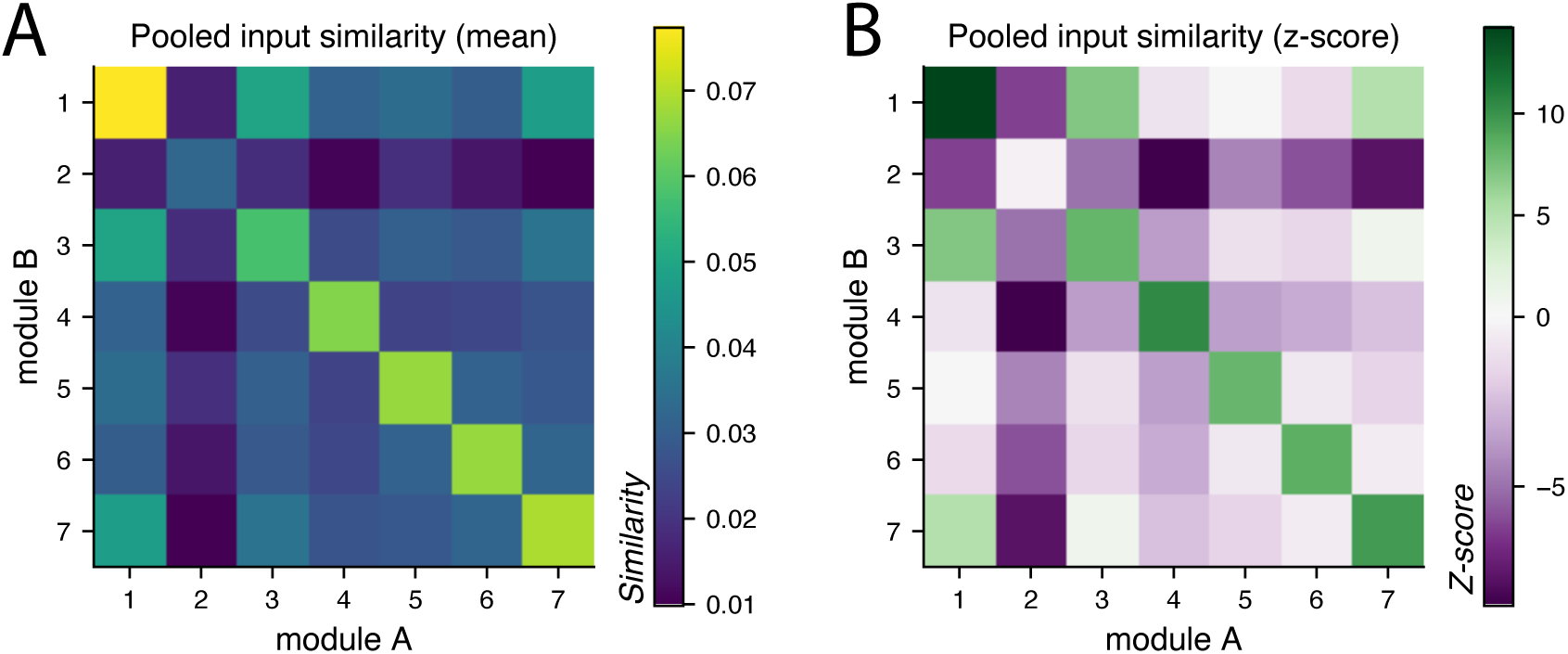
A. Mean pooled input similarity of the modules of the subconnectome. Subconnectome neurons have higher input similarity with neurons of the same module than with neurons from other modules (see diagonal). The inputs to neurons in the subconnectome of oviIN_R’s inputs include direct connections from neurons anywhere in the brain (except from neurons in the subconnectome). Heatmap colors indicate the mean input similarity between all neurons in module A vs. module B. B. Z-scores of pooled input similarity. Module labels were randomly shuffled (n = 100 shuffles) in a random model of input similarity. Along the diagonal, the true mean pooled input similarity was often several standard deviations away from that of the random model, indicating that the input self-similarity of the modules in the true clustering is not random. See full histograms in Figure 6-1.

## Discussion

With a central nervous system consisting of just under 140,000 neurons (***Dorkenwald et al., 2024***), the fruit fly, *Drosophila melanogaster*, exhibits the capacity for complex behavior driven by neuronal circuits that carry out sophisticated brain computations (***Scheffer and Meinertzhagen, 2021***; ***Griffith, 2013***; ***Oram and Card, 2022***; ***Haberkern and Jayaraman, 2016***). Context-dependent computations, which conditionally process or integrate inputs, endow an animal with behavioral flexibility (***Hindmarsh Sten et al., 2021***; ***Roemschied et al., 2023***; ***Devineni et al., 2019***). For instance, hungry flies persistently track a food odor despite repeated failure to find food, showing that need is selectively reinforced in such conditions (***Sayin et al., 2019***). Context can modulate egg-laying preference for acetic acid, despite the otherwise strong avoidance for this as food (***Joseph et al., 2009***). Additionally, circadian rhythms influence egg-laying, making oviposition more likely around the onset of night (***Cury et al., 2019***; ***Allemand, 1976***; ***Riva et al., 2024***). Processed sensory inputs integrate with context, memory, and state, in what we term high-level processing, to produce complex behavior.

The complexity inherent in the decision-making and behavior of *Drosophila* makes studying its tiny brain a worthwhile endeavor because isolating the circuits that drive high-level processing is within reach. The *Drosophila* brain has a hierarchical organization where peripheral sensory inputs undergo multiple stages of higher order processing, evident in both the adult (***Jeanne et al., 2018***; ***Kunin et al., 2023***) and larval brain (***Winding et al., 2023***; ***Betzel et al., 2024***). Researchers have made progress studying the sensory periphery and progressively discovering higher-order neurons and circuits (***Walker et al., 2024***; ***Jacobs et al., 2024***; ***Matsliah et al., 2024***), but such an approach is inadequate for discovering the circuits that drive the complex computations involved in contextdependent decision-making. Behaviors like oviposition depend on nearly every sensory modality, and starting at the periphery to discover the high-level circuits that integrate sensory inputs, state, and memory would require an expansive search. The connectome now offers another way to discover the neural circuits that are responsible for high-level processing. Although we have not identified verifiable circuits, our study takes a first step toward that goal.

The Hemibrain data has primarily been used to study well-characterized regions of the brain such as the CX (***Hulse et al., 2021***) and the MB (***Li et al., 2020***). The cell types and connectivity profiles of the CX circuits that compute abstract internal representations of angular orientation, maintained by self-motion signals and visual cues, have been cataloged by ***Hulse et al.*** (***2021***). Furthermore, they found that the output types of the CX project to various parts of the brain, including premotor and sensory areas, and that they also return to the CX, evoking a deep recurrent network. The MB supports associative learning, enabling flies to pair odorants with positive and negative experiences (***Handler et al., 2019***; ***Aso et al., 2014a***,b). Investigations using the connectome and in vivo calcium imaging have led to a detailed understanding of how the anterior paired lateral neuron (APL) normalizes Kenyon cell signals within the MB (***Prisco et al., 2021***). The connectome has also aided researchers in discovering new MB cell types that project outside of the MB lobes (***Li et al., 2020***) and that receive visual inputs as well as odor information (***Ganguly et al., 2024***; ***Li et al., 2020***). While great advances have been made in understanding these well-studied areas of the brain, there remains significant potential to leverage this dataset to investigate less well understood regions (***Scheffer and Meinertzhagen, 2021***; ***Lin et al., 2024***; ***Ito et al., 2014***).

In our approach, we analyzed the inputs to the oviposition circuit to study the neural correlates for the highest level computations that drive oviposition. We first observed that the oviIN is an outlier in the oviposition circuit and in the brain due to its abundant connectivity within and beyond the oviposition circuit. Given that the vast majority of inputs to oviIN are uncharacterized cell types, our goal was to identify the structural pathways into the oviposition circuit so that future work could determine the sensory and functional bases for these input pathways. We found that oviIN_R is the primary target of multiple cell types rather than solely collecting small amounts of input from neurons that primarily send outputs elsewhere. This pattern of connectivity is more aligned with a multi-circuit integrator role than a divisive normalizer. After applying a community-detection analysis, we found that the primary strong inputs to oviIN_R are distributed across multiple modules indicating that those primary strong inputs form the backbones of multiple distinct pathways to oviIN. Furthermore, we show that our modularity results do not reflect the same scope of anatomical organization that was found in the full Hemibrain modules by ***Kunin et al.*** (***2023***). Instead, maximizing modularity for the subconnectome containing oviIN_R’s inputs reveals an anatomical organization at a biophysically relevant scale for oviIN_R that is reminiscent of dendritic processing. Finally, we show that there is higher similarity among the inputs to neurons within a module than across modules, which further supports our interpretation of the modules as partitions around pathways that convey distinct information to oviIN.

Our study is limited to a partial brain connectome of a single specimen. Variability among individual female fruit flies may hinder the generality of our results, as well as the missing connections with neurons on the far left side of the central brain in the Hemibrain data. Additionally, the Hemibrain connectome only reconstructs classical chemical synaptic connections. Although electrical synapses and neuropeptides play an important role in the computations and functions of the *Drosophila* brain, gap junctions and neuropeptide release sites are not included in the current connectome data. Future iterations of *Drosophila* connectomes may include these additional features, enabling us to build upon the results we present here.

### High-level processing in the SNP

The majority of oviIN’s inputs come from the SNP -– an understudied region that includes the SLP, SIP, and SMP neuropils and that lacks a topological organization (***Ito et al., 2014***). These unstructured regions contain the components of circuits related to oviposition (***Zhang et al., 2020***; ***Wang et al., 2020***; ***Vijayan et al., 2023***), circadian rhythms (***Shafer et al., 2022***; ***Kaneko and Hall, 2000***; ***Fernández et al., 2008***; ***Reinhard et al., 2024***), olfaction (***Wang et al., 2014***; ***Ruta et al., 2010***; ***Aso et al., 2014a***; ***Cohn et al., 2015***; ***Séjourné et al., 2011***), and social behaviors like courtship and aggression (***Schretter et al., 2020***; ***Deutsch et al., 2020***; ***Coleman et al., 2024***; ***Taisz et al., 2023***; ***Wang et al., 2021***; ***Zhou et al., 2014***). The diffuse neuropils of the Protocerebrum are rife with high-level processing potential. MBONs carry odor information and send projections to the SNP at convergence zones where they meet the dendrites of dopaminergic neurons which respond to the valence of a stimulus (***Aso et al., 2014a***,b). Thus, the SNP are likely sites for circuits that are responsible for classical conditioning. Sensory and interoceptive inputs to the subesophageal zone are relayed to the SMP where they regulate water and food seeking behavior (***Landayan et al., 2021***; ***González Segarra et al., 2023***; ***Jourjine et al., 2016***). Taste projection neurons that project to the SLP are essential for conditioned taste aversion whereas those that bypass the SLP are not (***Kim et al., 2017***). The sexually dimorphic pC1 neurons have a role in persistent states related to courtship and aggression (***Deutsch et al., 2020***; ***Zhou et al., 2014***). They are presynaptic to oviIN, and according to the Hemibrain, they make most of their synapses in the SNP as well. Despite these behaviorally relevant functions, the SNP remain *fiterra incognitafi* as far as its organization (***Scheffer et al., 2020***; ***Ito et al., 2014***).

Without a pre-existing map of the SNP or a comprehensive wiring logic, we used modularity maximization to get a first approximation of the organization inherent in the abundant inputs to oviIN. To be clear, we do not claim to have identified complete (nor partial) circuits since we intentionally limited our modularity maximization to the network of neurons that are directly presynaptic to oviIN_R. Rather, our coarse clustering cast a wide net around the primary strong inputs to oviIN to dredge up clues about the regions and circuits that are associated with its inputs. Being able to associate uncharacterized cell types with each other, or with known cell types, is a far cry from discovering novel circuits, but it is a reasonable first step given that the SNP form abundant reciprocal connections among themselves and with other neuropils (***Lin et al., 2024***). This approach enabled us to determine to what extent the primary strong inputs are independent fispokesfi of a hub. Instead of the primary strong inputs coalescing into a single module, we found that there are primary strong inputs in every module. This suggests that there are distinct streams of inputs converging onto the oviIN. Ultimately, our approach revealed some approximate organization among oviIN’s inputs that may support the discovery of the circuits that are involved in the integration of sensory information, and the computations of evaluation and decision-making in oviposition. Future work will be needed to uncover and characterize the potential circuits that our connectivity analysis alludes to.

We observed that the FS1A neurons form recurrent connections among themselves and with other FB types, and they project to a recurrent network of SMP neurons that includes SMP386, SMP153, and SMP007. The FS1A neurons also project directly and strongly to oviIN. Thus we found two pathways along which FB information may flow to oviIN: a direct pathway via FS1A, and an indirect pathway from FS1A through a recurrently connected network of uncharacterized SMP neurons which themselves make strong, direct connections to oviIN. This suggests that the recurrently connected SMP neurons that are targeted by FS1A perform a distinct computation on the FB information received. Furthermore, since these SMP neurons also form recurrent connections with oviIN, they may be involved in a computation of FB input that is itself modulated by oviIN. It is possible that the SNP are organized into recurrently connected microcircuits such as this that perform high-level computations.

### Selective application of modularity analysis for structural scope

Network community detection methods generally fall into two classes: spectral methods which cluster nodes by leveraging the spectral properties of the graph to embed the network into a lowdimensional space (***Chung et al., 2021***), and modularity-based methods which partition nodes to maximize the difference between observed intra-cluster connections and those expected under a random graph model.

We used a modularity-maximization approach using the RenEEL machine learning scheme (***Guo et al., 2019***) because of its ease of interpretation. Its emphasis on intra-connectedness of the node communities (***Newman, 2006***; ***Newman and Girvan, 2004***; ***Guo et al., 2023***; ***Kunin et al., 2023***) reveals assortative groups of neurons and cell types. By forming densely connected modules within the subconnectome of oviIN_R’s inputs, our modularity-based method offers insight into the structural organization of this network and provides predictions for the functions of the cell types that are clustered together.

In our study, we chose to apply RenEEL to a subnetwork of the connectome to control the scope of the structure we were probing. RenEEL was previously applied to the full Hemibrain connectome by ***Kunin et al.*** (***2023***) and had recovered gross anatomical structure that reflects the neuropil organization of the brain, as well as finer structure such as the layers of the FB. While such macroscale anatomical structure often underlies functional circuit structure (***Betzel et al., 2013***), it might obscure mesoscale and microscale circuit structure that is not necessarily contained within a given neuropil (***Betzel et al., 2018***; ***Betzel, 2023***). For example, the FS1A and the SMP386 neurons are from different neuropils and they are clustered into separate modules for the full Hemibrain connectome; however, they are clustered into the same module when RenEEL is applied to the subconnectome of oviIN_R’s inputs. These neurons are separately partitioned in the full connectome because the connections they make with other neurons from their own neuropil are denser than any connections made with neurons outside of their neuropils. When the scope of the network is limited only to neurons that are involved in a particular function, such as oviposition, the interactions between these neurons becomes more apparent. By applying RenEEL to a subconnectome consisting of the inputs to oviIN_R, we limited our analysis to finding partitions at a mesoscale and microscale level of relevance.

Generalized modularity [eqn.1] offers an established method with which to probe mesoscale and microscale structure. The resolution parameter can be adjusted to retrieve more modules containing fewer neurons. A previous study demonstrated that maximizing generalized modularity reveals the hierarchically nested anatomical structure of the brain when applied to the full Hemibrain network (***Kunin et al., 2023***). The modules identified at higher resolutions were generally subsets of the coarsest modules, though this partitioning was not always strict. Even still, the high resolution modules of the full Hemibrain did not correspond to the coarse modules that we obtained for the subconnectome consisting of the inputs to oviIN_R (***Figure 5***B,C). This demonstrates that maximizing classical (i.e. coarse) modularity on a subconnectome does not produce the same results as maximizing generalized modularity at higher resolutions on a full connectome. Although our study did not set out to investigate the subtleties between classical and generalized modularity analyses, we propose that applying classical modularity to a subconnectome targets a different scope of brain structure than applying generalized modularity on the full connectome. Future work is needed to gain a comprehensive understanding of how community-detection tools can be honed to reliably retrieve circuit structure in the *Drosophila* brain.

## Acknowledgments

We would like to thank Stephen Thornquist for his involvement in the early part of this study, and Maria de la Paz Fernandez, Larry Abbott, and Ashok Litwin-Kumar for their helpful feedback. GJG and RAWL were supported by NIH NINDS grant K22NS104187.

## Extended Data

**Figure 1-1.**
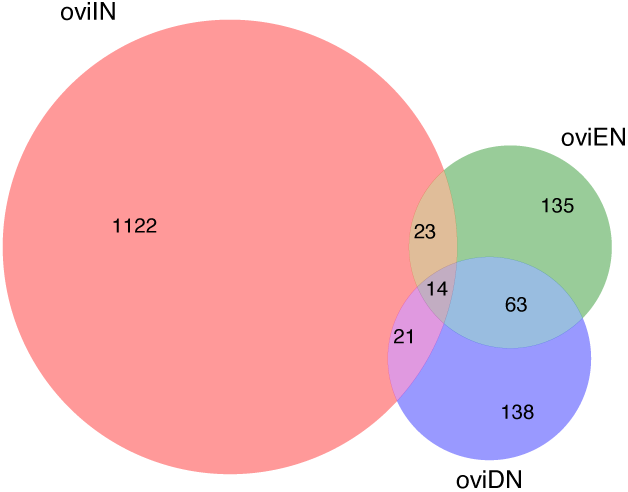
The overlap in unique presynaptic partners to oviIN_R (pink), SMP550_R (oviEN; green), and the combined group of right oviDN subtypes (oviDNa_R, oviDNb_R, and both instances of SLP410_R; purple). There are 14 unique neurons that are presynaptic to all 3 cell types.

**Table 4-1.** Cell types in each module of the subconnectome of oviIN_R’s inputs. The module is indicated in the second column. The third column contains the total synapse weight onto oviIN_R by the neurons of the indicated cell type in the indicated module.

**Figure 4-1a.**
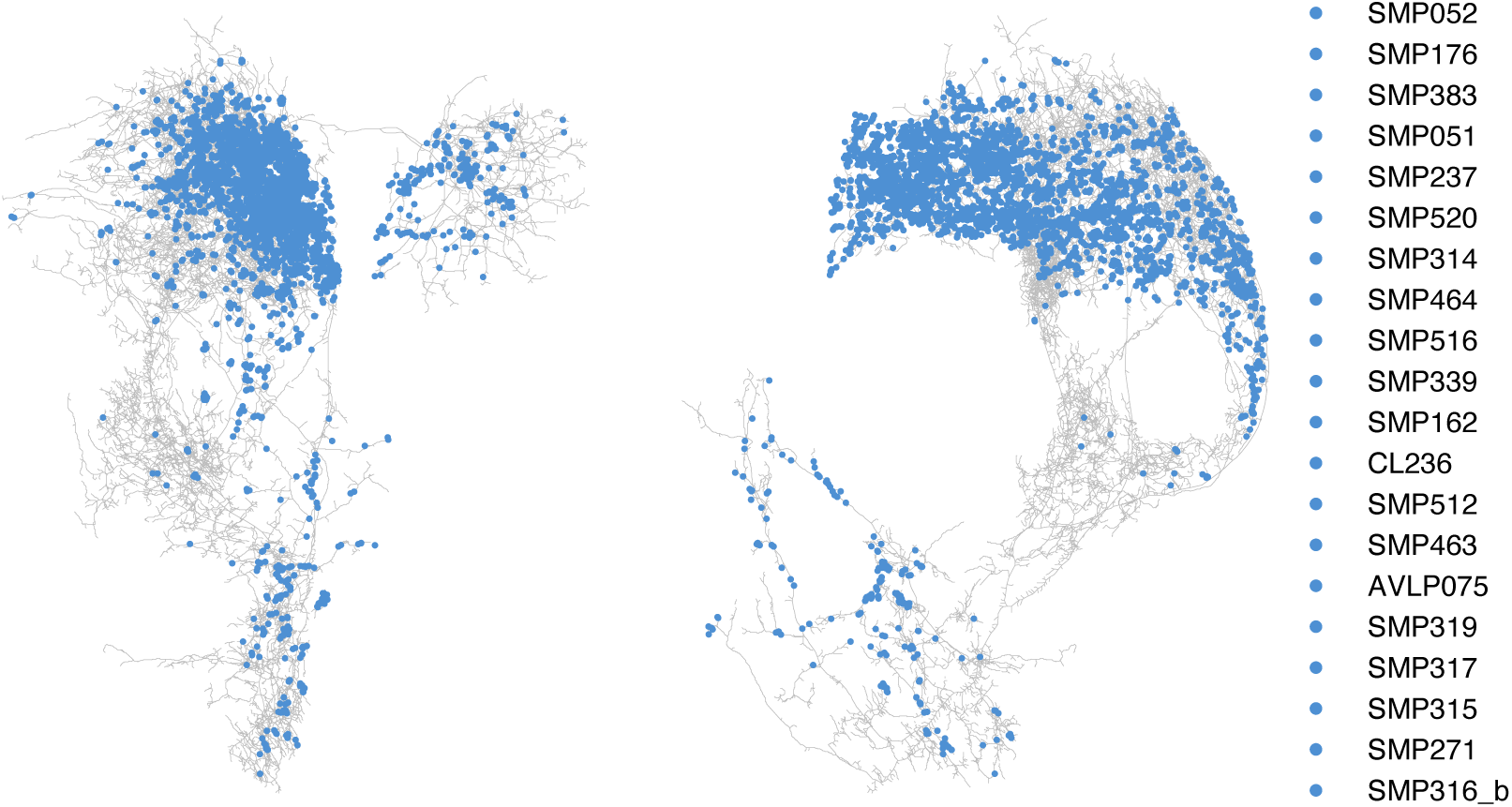
Synaptic input sites on oviIN_R from module 1. Left panel: front view; Right panel: lateral view. The 20 cell types from module 1 making the strongest connections to oviIN_R are shown in the legend.

**Figure 4-1b.**
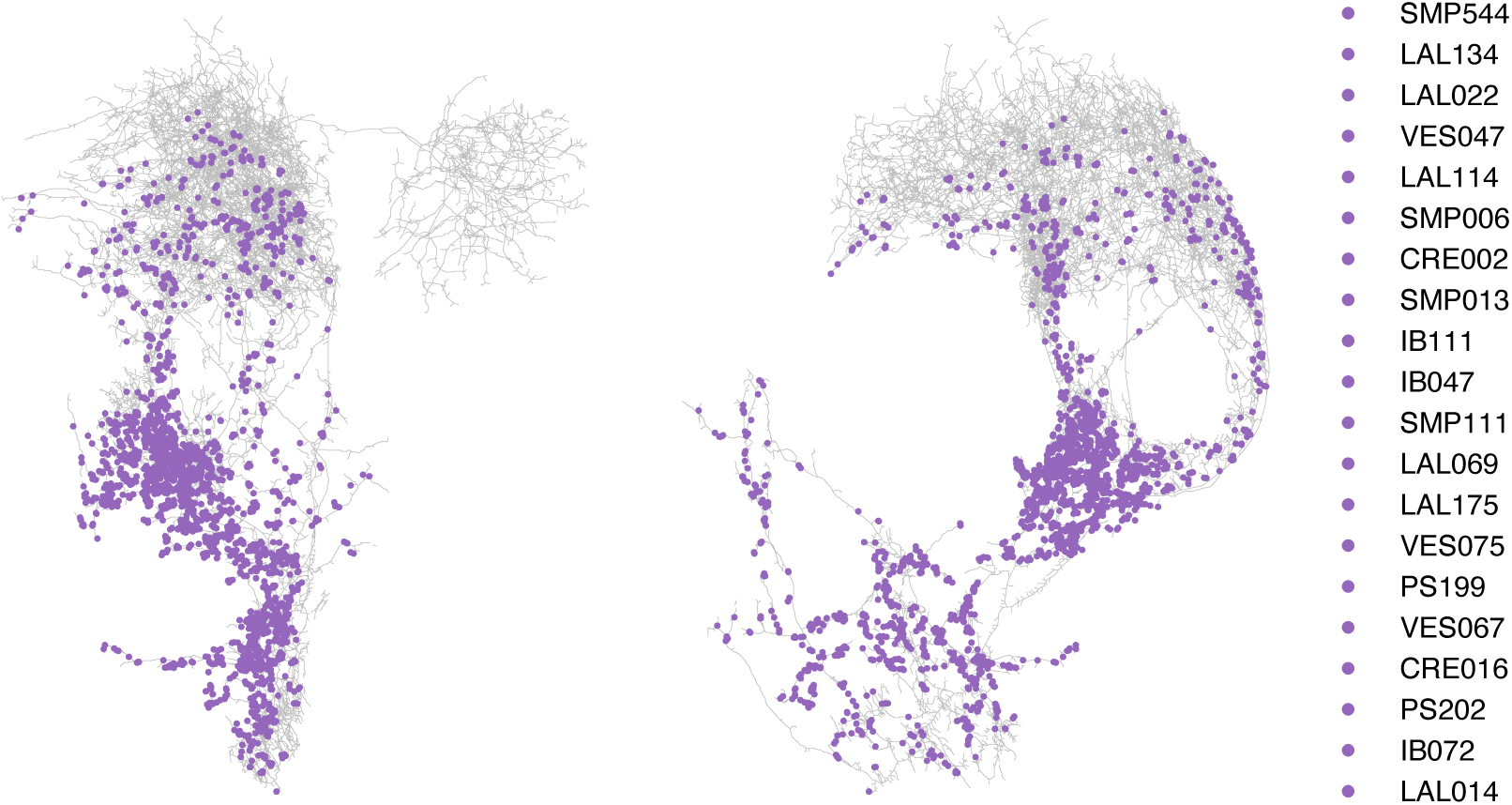
Synaptic input sites on oviIN_R from module 2. Left panel: front view; Right panel: lateral view. The 20 cell types from module 2 making the strongest connections to oviIN_R are shown in the legend.

**Figure 4-1c.**
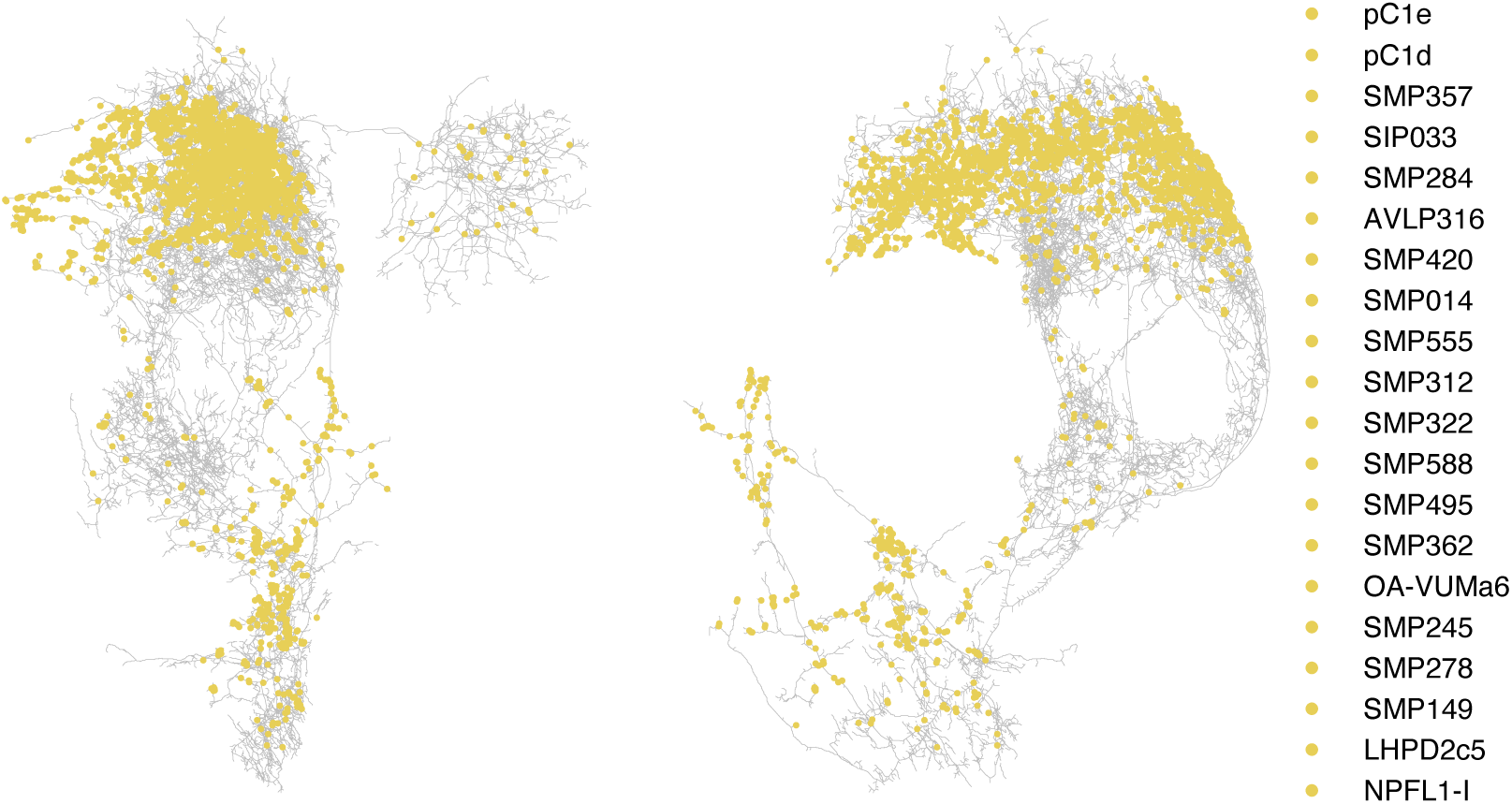
Synaptic input sites on oviIN_R from module 3. Left panel: front view; Right panel: lateral view. The 20 cell types from module 3 making the strongest connections to oviIN_R are shown in the legend.

**Figure 4-1d.**
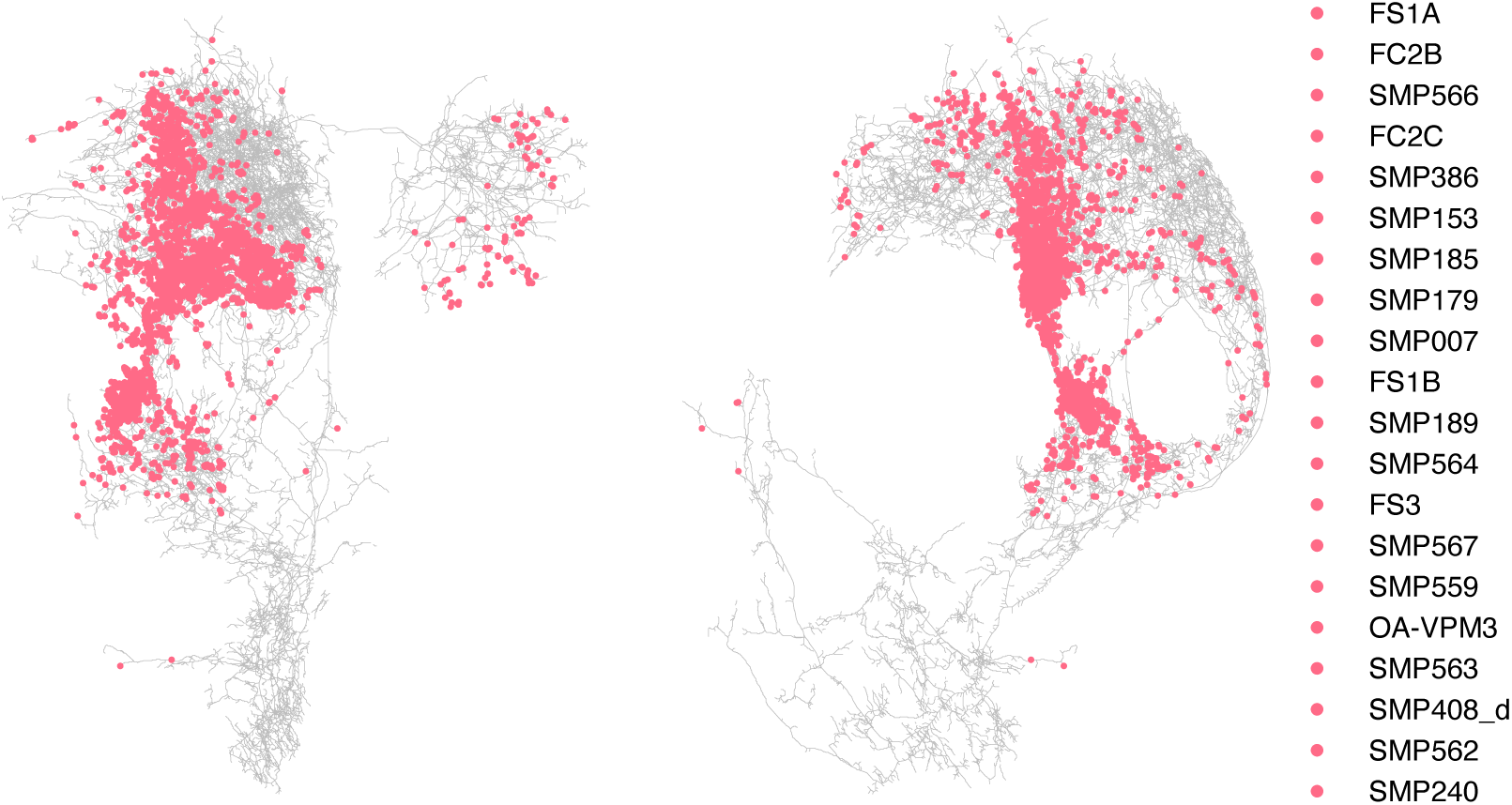
Synaptic input sites on oviIN_R from module 4. Left panel: front view; Right panel: lateral view. The 20 cell types from module 4 making the strongest connections to oviIN_R are shown in the legend.

**Figure 4-1e.**
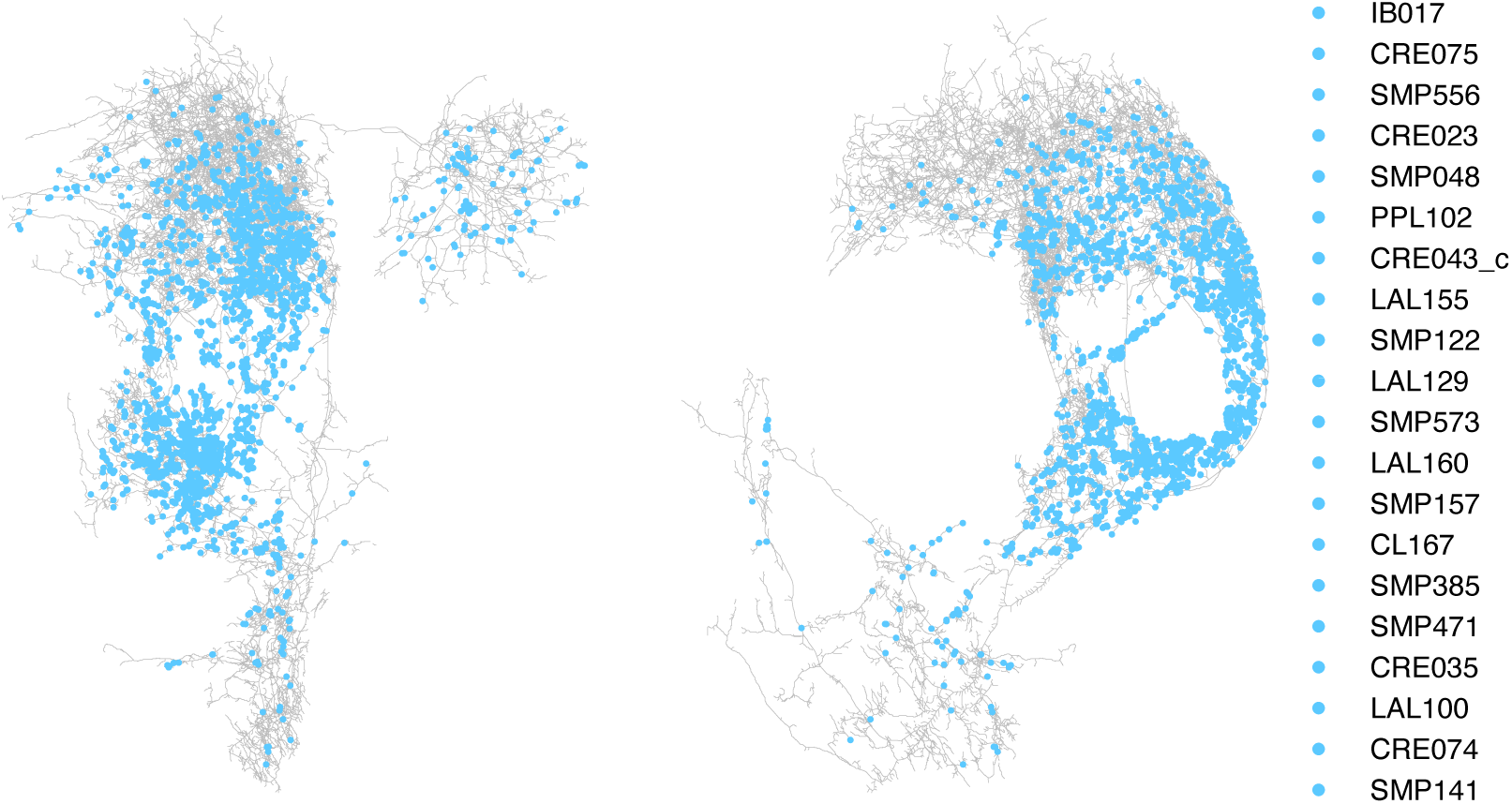
Synaptic input sites on oviIN_R from module 5. Left panel: front view; Right panel: lateral view. The 20 cell types from module 5 making the strongest connections to oviIN_R are shown in the legend.

**Figure 4-1f.**
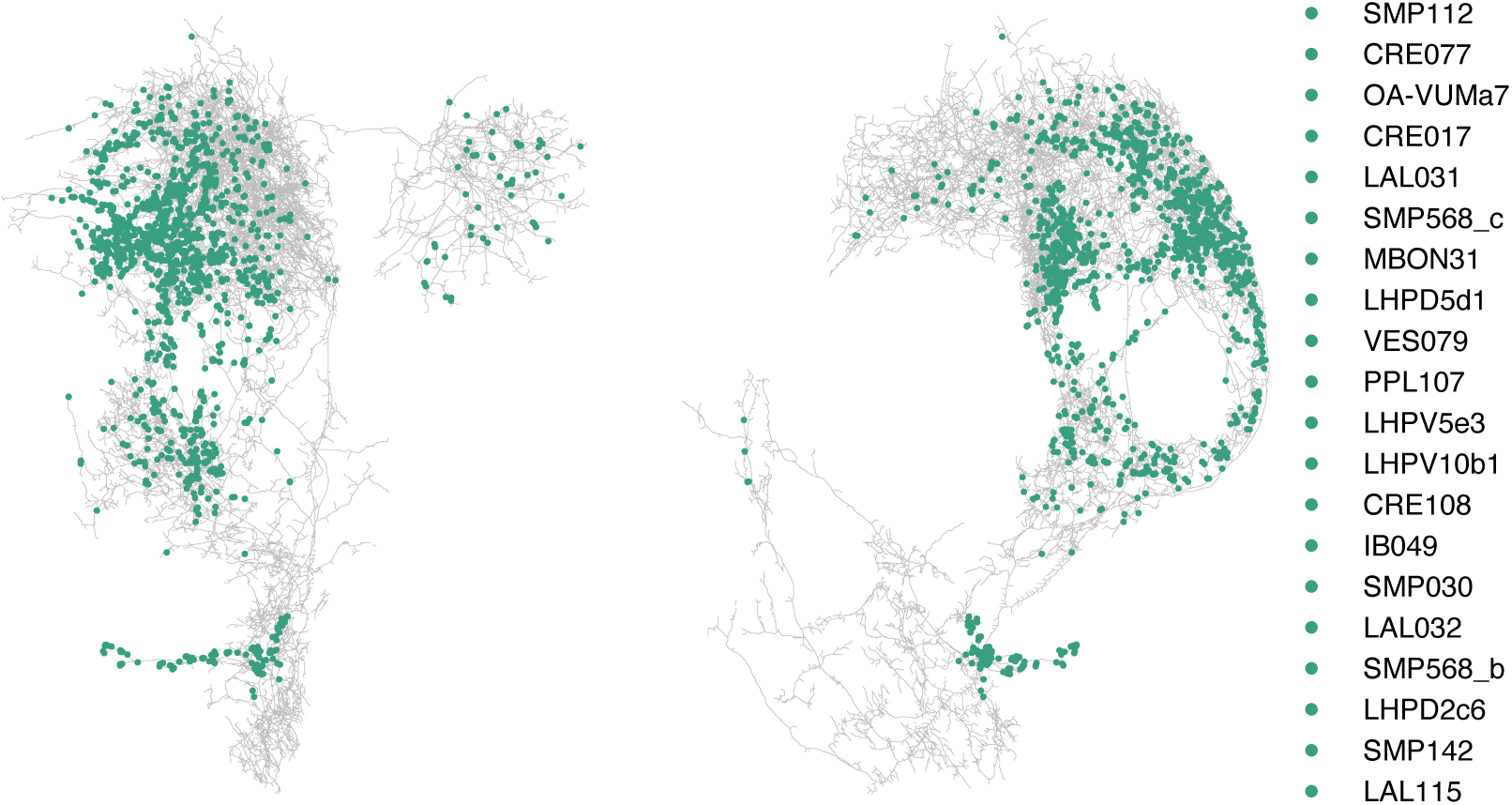
Synaptic input sites on oviIN_R from module 6. Left panel: front view; Right panel: lateral view. The 20 cell types from module 6 making the strongest connections to oviIN_R are shown in the legend.

**Figure 4-1g.**
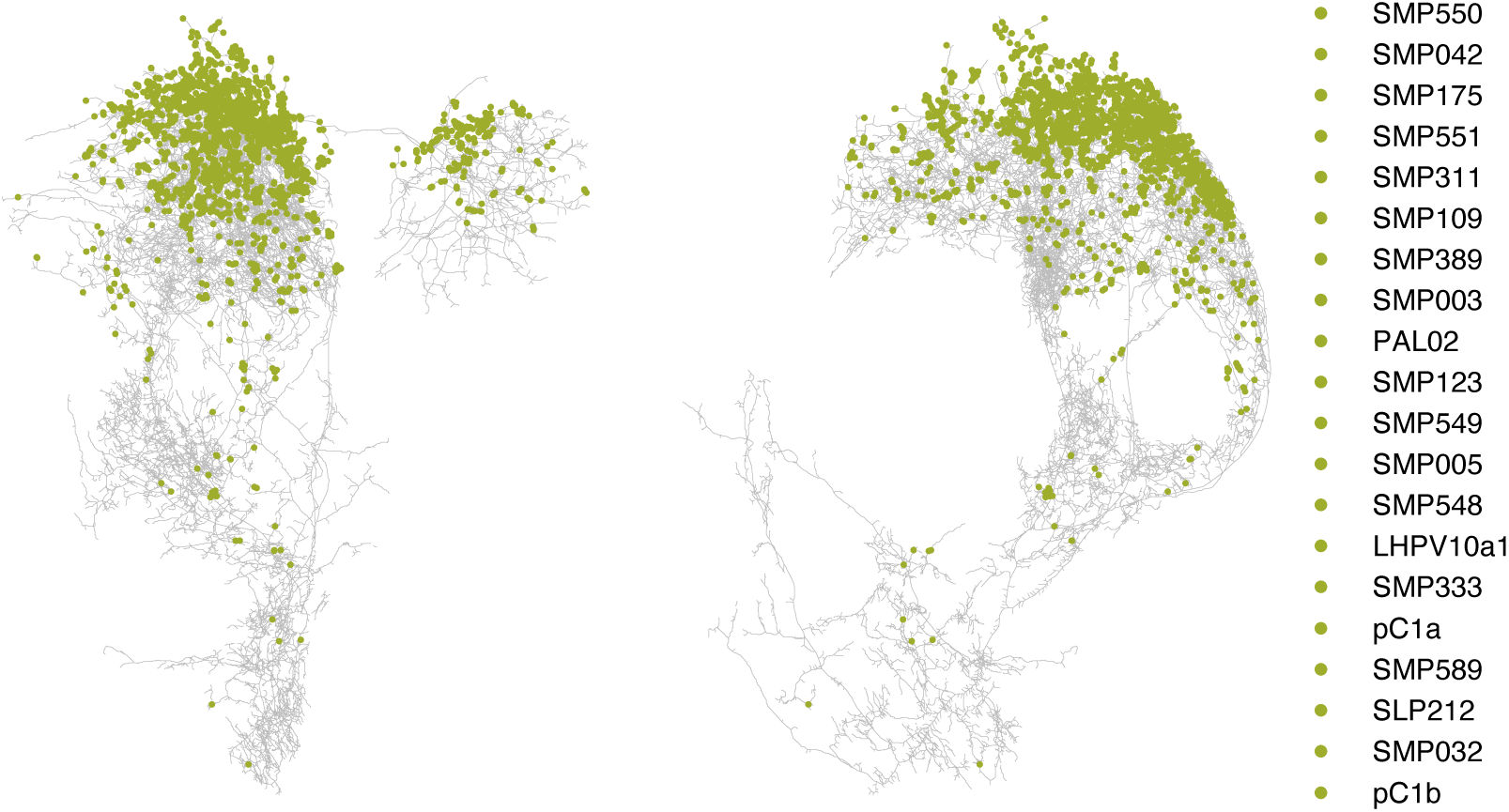
Synaptic input sites on oviIN_R from module 7. Left panel: front view; Right panel: lateral view. The 20 cell types from module 7 making the strongest connections to oviIN_R are shown in the legend.

**Figure 4-2.**
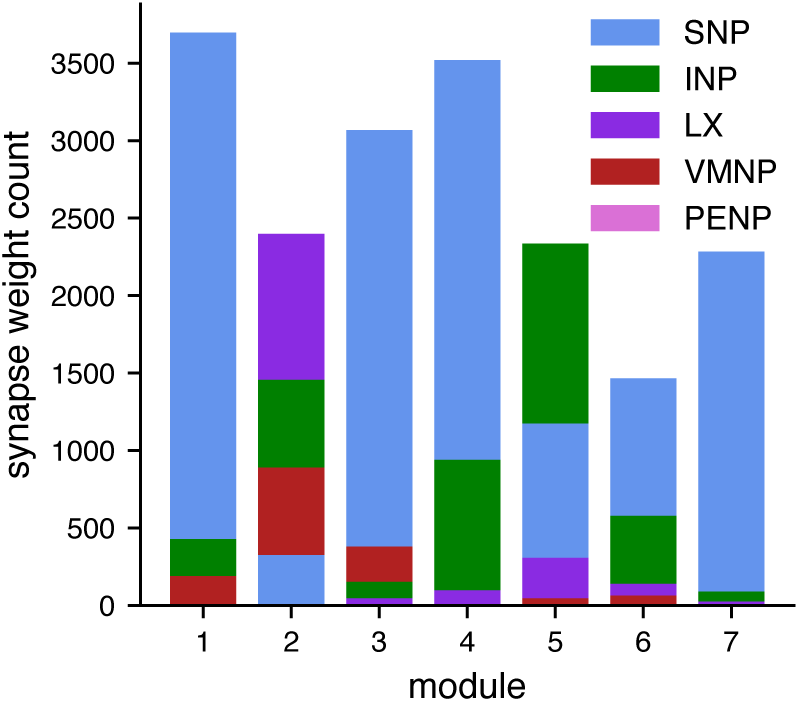
Supercategories where synapses from presynaptic partners to oviIN_R are localized. For the neurons within each module, the counts of synapses to oviIN_R within a neuropil are used to compute the supercategory synapse weight counts (see ***Figure 5-1***).

**Figure 5-1.**
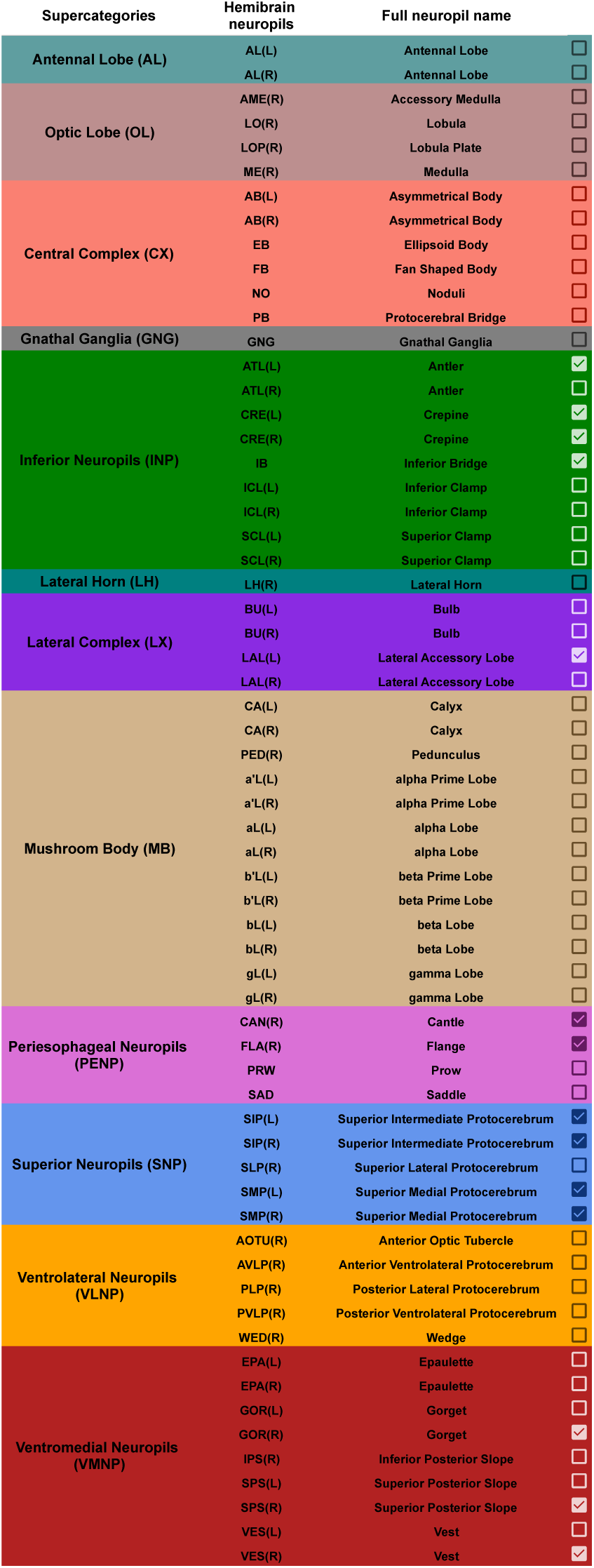
All Hemibrain neuropils from the v1.2.1 dataset along with their supercategories as described by ***Ito et al.*** (***2014***). The colors of each row correspond to the supercategory colors used in ***Figure 5***A and ***Figure 4-2***. A check mark in the last column denotes a neuropil where the oviIN_R receives connections.

**Figure 6-1.**
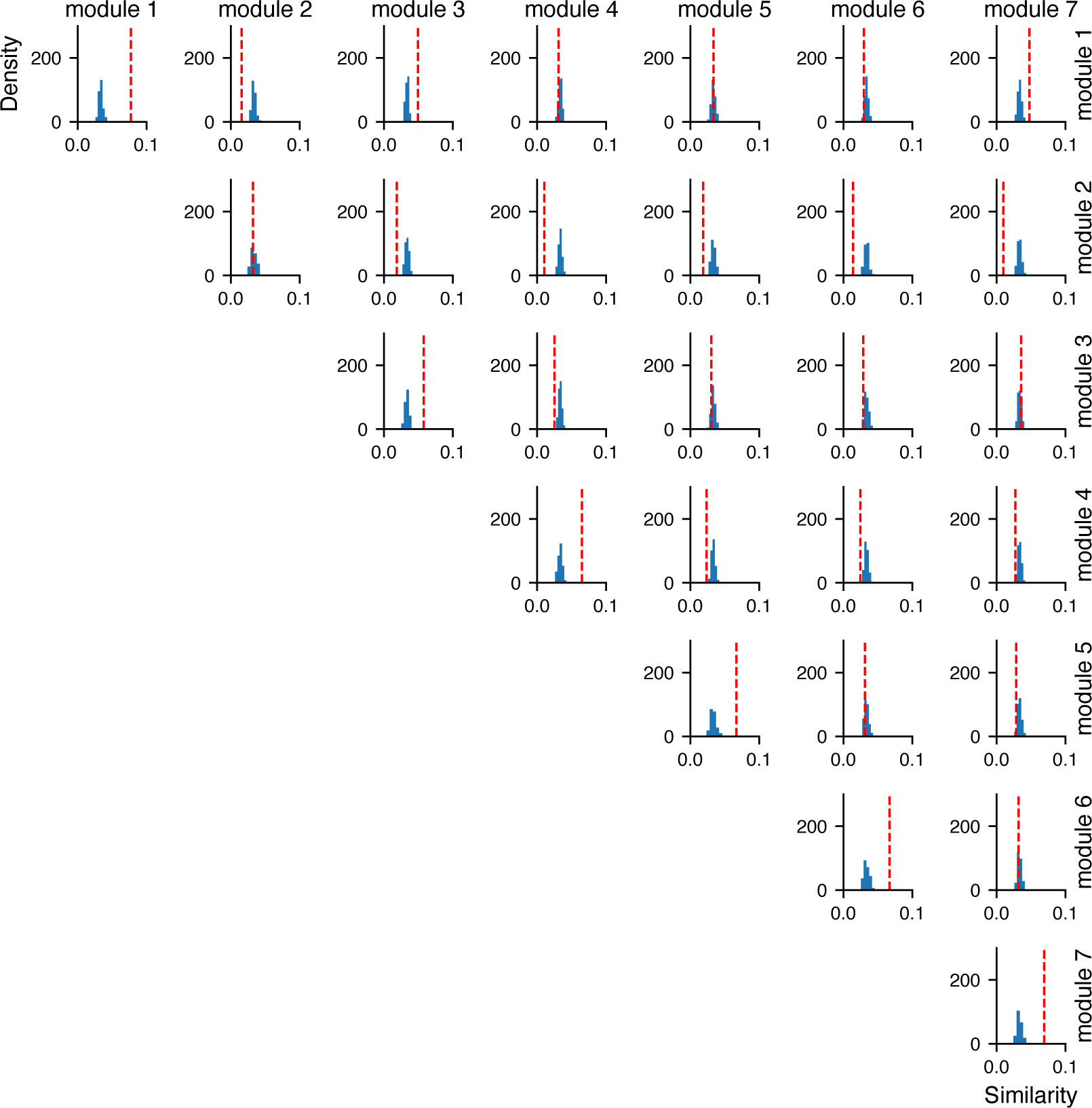
Randomization of module labels (n = 100 shuffles) shows decreased similarity for self-same cluster pairs. Input similarity is shown on the *x*-axes (similarity ranges from 0 to 1). The counts of similarity values are shown on the *y*-axes. The red dashed line in each plot indicates the mean input similarity between modules as visualized in ***Figure 6***A. The histograms show the similarity scores for the random models. Along the diagonal, the mean input similarity among neurons in the same module (red dashed line) tends to be higher than the mean input similarity among neurons from random modules. In off-diagonal plots, the mean input similarity between neurons from two different modules (red dashed line) tends to be within the similarity distribution for neurons from random modules or lower than the distribution mean.

## References

Allemand R. Les rythmes de vitellogenese et d’ovulation en photoperiode LD 12:12 de *Drosophila melanogaster*. Journal of Insect Physiology. 1976 Jan; 22(7):1031–1035. https://www.sciencedirect.com/science/article/pii/0022191076900883, doi: 10.1016/0022-1910(76)90088-3.

Aso Y, Hattori D, Yu Y, Johnston RM, Iyer NA, Ngo TT, Dionne H, Abbott L, Axel R, Tanimoto H, Rubin GM. The neuronal architecture of the mushroom body provides a logic for associative learning. eLife. 2014 Dec; 3:e04577. 10.7554/eLife.04577, doi: 10.7554/eLife.04577.

Aso Y, Sitaraman D, Ichinose T, Kaun KR, Vogt K, Belliart-Guérin G, Plaçais PY, Robie AA, Yamagata N, Schnaitmann C, Rowell WJ, Johnston RM, Ngo TTB, Chen N, Korff W, Nitabach MN, Heberlein U, Preat T, Branson KM, Tanimoto H, et al. Mushroom body output neurons encode valence and guide memory-based action selection in Drosophila. eLife. 2014 Dec; 3:e04580. 10.7554/eLife.04580, doi: 10.7554/eLife.04580.

Azanchi R, Kaun KR, Heberlein U. Competing dopamine neurons drive oviposition choice for ethanol in Drosophila. Proceedings of the National Academy of Sciences. 2013 Dec; 110(52):21153–21158. https://www.pnas.org/doi/full/10.1073/pnas.1320208110, doi: 10.1073/pnas.1320208110.

Bailly TPM, Kohlmeier P, Etienne RS, Wertheim B, Billeter JC. Social modulation of oogenesis and egg laying in Drosophila melanogaster. Current Biology. 2023 Jul; 33(14):2865–2877.e4. https://www.cell.com/current-biology/abstract/S0960-9822(23)00750-9, doi: 10.1016/j.cub.2023.05.074, publisher: Elsevier.

Betzel R, Puxeddu MG, Seguin C. Hierarchical communities in the larval Drosophila connectome: Links to cellular annotations and network topology. Proceedings of the National Academy of Sciences. 2024 Sep; 121(38):e2320177121. https://www.pnas.org/doi/abs/10.1073/pnas.2320177121, doi: 10.1073/pnas.2320177121, publisher: Proceedings of the National Academy of Sciences.

Betzel RF. Chapter 7 - Community detection in network neuroscience. In: Schirmer MD, Arichi T, Chung AW, editors. Connectome Analysis Academic Press; 2023.p. 149–171. https://www.sciencedirect.com/science/article/pii/B9780323852807000166, doi: 10.1016/B978-0-323-85280-7.00016-6.

Betzel RF, Griffa A, Avena-Koenigsberger A, Goñi J, Thiran JP, Hagmann P, Sporns O. Multi-scale community organization of the human structural connectome and its relationship with resting-state functional connectivity. Network Science. 2013 Dec; 1(3):353–373. https://www.cambridge.org/core/journals/network-science/article/multiscale-community-organization-of-the-human-structural-connectome-and-its-relationship-with-restingstate-functional-co97B9B699236A2F129684F1D80D1DA220, doi: 10.1017/nws.2013.19.

Betzel RF, Medaglia JD, Bassett DS. Diversity of meso-scale architecture in human and non-human connectomes. Nature Communications. 2018 Jan; 9(1):346. https://www.nature.com/articles/s41467-017-02681-z, doi: 10.1038/s41467-017-02681-z, publisher: Nature Publishing Group.

Chou YH, Spletter ML, Yaksi E, Leong JCS, Wilson RI, Luo L. Diversity and wiring variability of olfactory local interneurons in the Drosophila antennal lobe. Nature Neuroscience. 2010 Apr; 13(4):439–449. https://www.nature.com/articles/nn.2489, doi: 10.1038/nn.2489, publisher: Nature Publishing Group.

Chung J, Bridgeford E, Arroyo J, Pedigo BD, Saad-Eldin A, Gopalakrishnan V, Xiang L, Priebe CE, Vogelstein JT. Statistical Connectomics. Annual Review of Statistics and Its Application. 2021 Mar; 8(Volume 8, 2021):463–492. https://www.annualreviews.org/content/journals/10.1146/annurev-statistics-042720-023234, doi: 10.1146/annurev-statistics-042720-023234, publisher: Annual Reviews.

Churchill ER, Dytham C, Bridle JR, Thom MDF. Social and physical environment independently affect oviposition decisions in Drosophila. Behavioral Ecology. 2021 Nov; 32(6):1391–1399. 10.1093/beheco/arab105, doi: 10.1093/beheco/arab105.

Cohn R, Morantte I, Ruta V. Coordinated and Compartmentalized Neuromodulation Shapes Sensory Processing in Drosophila. Cell. 2015 Dec; 163(7):1742–1755. https://www.cell.com/cell/abstract/S0092-8674(15)01499-3, doi: 10.1016/j.cell.2015.11.019, publisher: Elsevier.

Coleman RT, Morantte I, Koreman GT, Cheng ML, Ding Y, Ruta V. A modular circuit coordinates the diversification of courtship strategies. Nature. 2024 Oct; p. 1–9. https://www.nature.com/articles/s41586-024-08028-1, doi: 10.1038/s41586-024-08028-1, publisher: Nature Publishing Group.

Cury KM, Axel R. Flexible neural control of transition points within the egg-laying behavioral sequence in Drosophila. Nature Neuroscience. 2023 Jun; 26(6):1054–1067. https://www.nature.com/articles/s41593-023-01332-5, doi: 10.1038/s41593-023-01332-5, number: 6 Publisher: Nature Publishing Group.

Cury KM, Prud’homme B, Gompel N. A short guide to insect oviposition: when, where and how to lay an egg. Journal of Neurogenetics. 2019 Apr; 33(2):75–89. 10.1080/01677063.2019.1586898, doi: 10.1080/01677063.2019.1586898, publisher: Taylor & Francis _eprint: https://doi.org/10.1080/01677063.2019.1586898.

Deutsch D, Pacheco D, Encarnacion-Rivera L, Pereira T, Fathy R, Clemens J, Girardin C, Calhoun A, Ireland E, Burke A, Dorkenwald S, McKellar C, Macrina T, Lu R, Lee K, Kemnitz N, Ih D, Castro M, Halageri A, Jordan C, et al. The neural basis for a persistent internal state in Drosophila females. eLife. 2020 Nov; 9:e59502. 10.7554/eLife.59502, doi: 10.7554/eLife.59502.

Devineni AV, Sun B, Zhukovskaya A, Axel R. Acetic acid activates distinct taste pathways in Drosophila to elicit opposing, state-dependent feeding responses. eLife. 2019 Jun; 8:e47677. 10.7554/eLife.47677, doi: 10.7554/eLife.47677, publisher: eLife Sciences Publications, Ltd.

Dorkenwald S, Matsliah A, Sterling AR, Schlegel P, Yu Sc, McKellar CE, Lin A, Costa M, Eichler K, Yin Y, Silversmith W, Schneider-Mizell C, Jordan CS, Brittain D, Halageri A, Kuehner K, Ogedengbe O, Morey R, Gager J, Kruk K, et al. Neuronal wiring diagram of an adult brain. Nature. 2024 Oct; 634(8032):124–138. https://www.nature.com/articles/s41586-024-07558-y, doi: 10.1038/s41586-024-07558-y, publisher: Nature Publishing Group.

Feng K, Palfreyman MT, Häsemeyer M, Talsma A, Dickson BJ. Ascending SAG Neurons Control Sexual Receptivity of Drosophila Females. Neuron. 2014 Jul; 83(1):135–148. https://www.sciencedirect.com/science/article/pii/S0896627314004036, doi: 10.1016/j.neuron.2014.05.017.

Fernández MP, Berni J, Ceriani MF. Circadian Remodeling of Neuronal Circuits Involved in Rhythmic Behavior. PLOS Biology. 2008 Mar; 6(3):e69. https://journals.plos.org/plosbiology/article?id=10.1371/journal.pbio.0060069, doi: 10.1371/journal.pbio.0060069.

Ganguly I, Heckman EL, Litwin-Kumar A, Clowney EJ, Behnia R. Diversity of visual inputs to Kenyon cells of the Drosophila mushroom body. Nature Communications. 2024 Jul; 15(1):5698. https://www.nature.com/articles/s41467-024-49616-z, doi: 10.1038/s41467-024-49616-z, publisher: Nature Publishing Group.

González Segarra AJ, Pontes G, Jourjine N, Del Toro A, Scott K. Hunger- and thirst-sensing neurons modulate a neuroendocrine network to coordinate sugar and water ingestion. eLife. 2023 Sep; 12:RP88143. 10.7554/eLife.88143, doi: 10.7554/eLife.88143, publisher: eLife Sciences Publications, Ltd.

Griffith LC. Neuromodulatory control of sleep in Drosophila melanogaster: integration of competing and complementary behaviors. Current Opinion in Neurobiology. 2013 Oct; 23(5):819–823. https://linkinghub.elsevier.com/retrieve/pii/S0959438813001165, doi: 10.1016/j.conb.2013.05.003.

Guo J, Singh P, Bassler KE. Reduced network extremal ensemble learning (RenEEL) scheme for community detection in complex networks. Scientific Reports. 2019 Oct; 9(1):14234. https://www.nature.com/articles/s41598-019-50739-3, doi: 10.1038/s41598-019-50739-3, number: 1 Publisher: Nature Publishing Group.

Guo J, Singh P, Bassler KE. Resolution limit revisited: community detection using generalized modularity density. Journal of Physics: Complexity. 2023 Mar; 4(2):025001. 10.1088/2632-072X/acc4a4, doi: 10.1088/2632-072X/acc4a4, publisher: IOP Publishing.

Haberkern H, Jayaraman V. Studying small brains to understand the building blocks of cognition. Current Opinion in Neurobiology. 2016 Apr; 37:59–65. https://www.sciencedirect.com/science/article/pii/S0959438816000088, doi: 10.1016/j.conb.2016.01.007.

Handler A, Graham TGW, Cohn R, Morantte I, Siliciano AF, Zeng J, Li Y, Ruta V. Distinct Dopamine Receptor Pathways Underlie the Temporal Sensitivity of Associative Learning. Cell. 2019 Jun; 178(1):60–75.e19. https://www.sciencedirect.com/science/article/pii/S0092867419306117, doi: 10.1016/j.cell.2019.05.040.

Haynes PR, Christmann BL, Griffith LC. A single pair of neurons links sleep to memory consolidation in Drosophila melanogaster. eLife. 2015 Jan; 4:e03868. 10.7554/eLife.03868, doi: 10.7554/eLife.03868.

Helfrich-Förster C, Shafer OT, Wülbeck C, Grieshaber E, Rieger D, Taghert P. Development and morphology of the clock-gene-expressing lateral neurons of Drosophila melanogaster. Journal of Comparative Neurology. 2007; 500(1):47–70. https://onlinelibrary.wiley.com/doi/abs/10.1002/cne.21146, doi: 10.1002/cne.21146, _eprint: https://onlinelibrary.wiley.com/doi/pdf/10.1002/cne.21146.

Hindmarsh Sten T, Li R, Otopalik A, Ruta V. Sexual arousal gates visual processing during Drosophila courtship. Nature. 2021 Jul; 595(7868):549–553. https://www.nature.com/articles/s41586-021-03714-w, doi: 10.1038/s41586-021-03714-w, publisher: Nature Publishing Group.

Hulse BK, Haberkern H, Franconville R, Turner-Evans DB, Takemura Sy, Wolff T, Noorman M, Dreher M, Dan C, Parekh R, Hermundstad AM, Rubin GM, Jayaraman V. A connectome of the Drosophila central complex reveals network motifs suitable for flexible navigation and context-dependent action selection. eLife. 2021 Oct; 10:e66039. https://elifesciences.org/articles/66039, doi: 10.7554/eLife.66039.

Imoto K, Ishikawa Y, Aso Y, Funke J, Tanaka R, Kamikouchi A. Neural-circuit basis of song preference learning in fruit flies. iScience. 2024 Jul; 27(7):110266. https://www.sciencedirect.com/science/article/pii/S2589004224014913, doi: 10.1016/j.isci.2024.110266.

Ito K, Shinomiya K, Ito M, Armstrong JD, Boyan G, Hartenstein V, Harzsch S, Heisenberg M, Homberg U, Jenett A, Keshishian H, Restifo LL, Rössler W, Simpson JH, Strausfeld NJ, Strauss R, Vosshall LB. A Systematic Nomenclature for the Insect Brain. Neuron. 2014 Feb; 81(4):755–765. https://www.sciencedirect.com/science/article/pii/S0896627313011781, doi: 10.1016/j.neuron.2013.12.017.

Jacobs RV, Wang CX, Nguyen L, Pruitt TJ, Wang P, Lozada-Perdomo FV, Deere JU, Liphart HA, Devineni AV. Overlap and divergence of neural circuits mediating distinct behavioral responses to sugar. Cell Reports. 2024 Oct; 43(10):114782. https://www.sciencedirect.com/science/article/pii/S2211124724011331, doi: 10.1016/j.celrep.2024.114782.

Jeanne JM, Fişek M, Wilson RI. The Organization of Projections from Olfactory Glomeruli onto Higher-Order Neurons. Neuron. 2018 Jun; 98(6):1198–1213.e6. https://www.cell.com/neuron/abstract/S0896-6273(18)30383-0, doi: 10.1016/j.neuron.2018.05.011.

Joseph RM, Devineni AV, King IFG, Heberlein U. Oviposition preference for and positional avoidance of acetic acid provide a model for competing behavioral drives in Drosophila. Proceedings of the National Academy of Sciences. 2009 Jul; 106(27):11352–11357. https://www.pnas.org/doi/full/10.1073/pnas.0901419106, doi: 10.1073/pnas.0901419106.

Jourjine N, Mullaney BC, Mann K, Scott K. Coupled Sensing of Hunger and Thirst Signals Balances Sugar and Water Consumption. Cell. 2016 Aug; 166(4):855–866. https://www.sciencedirect.com/science/article/pii/S0092867416308546, doi: 10.1016/j.cell.2016.06.046.

Kaneko M, Hall JC. Neuroanatomy of cells expressing clock genes in Drosophila: Transgenic manipulation of the period and timeless genes to mark the perikarya of circadian pacemaker neurons and their projections. Journal of Comparative Neurology. 2000; 422(1):66–94. https://onlinelibrary.wiley.com/doi/abs/10.1002/%28SICI%291096-9861%2820000619%29422%3A1%3C66%3A%3AAID-CNE5%3E3.0.CO%3B2-2, doi: 10.1002/(SICI)1096-9861(20000619)422:1<66::AID-CNE5>3.0.CO;2-2, _eprint: https://onlinelibrary.wiley.com/doi/pdf/10.1002/%28SICI%291096-9861%2820000619%29422%3A1%3C66%3A%3AAID-CNE5%3E3.0.CO%3B2-2.

Keene AC, Krashes MJ, Leung B, Bernard JA, Waddell S. *Drosophila* Dorsal Paired Medial Neurons Provide a General Mechanism for Memory Consolidation. Current Biology. 2006 Aug; 16(15):1524–1530. https://www.sciencedirect.com/science/article/pii/S0960982206017076, doi: 10.1016/j.cub.2006.06.022.

Keene AC, Stratmann M, Keller A, Perrat PN, Vosshall LB, Waddell S. Diverse Odor-Conditioned Memories Require Uniquely Timed Dorsal Paired Medial Neuron Output. Neuron. 2004 Oct; 44(3):521–533. https://www.sciencedirect.com/science/article/pii/S0896627304006476, doi: 10.1016/j.neuron.2004.10.006.

Kim H, Kirkhart C, Scott K. Long-range projection neurons in the taste circuit of Drosophila. eLife. 2017 Feb; 6:e23386. 10.7554/eLife.23386, doi: 10.7554/eLife.23386.

Kunin AB, Guo J, Bassler KE, Pitkow X, Josić K. Hierarchical Modular Structure of the Drosophila Connectome. Journal of Neuroscience. 2023 Sep; 43(37):6384–6400. https://www.jneurosci.org/content/43/37/6384, doi: 10.1523/JNEUROSCI.0134-23.2023, publisher: Society for Neuroscience Section: Research Articles.

Landayan D, Wang BP, Zhou J, Wolf FW. Thirst interneurons that promote water seeking and limit feeding behavior in Drosophila. eLife. 2021 May; 10:e66286. https://www.ncbi.nlm.nih.gov/pmc/articles/PMC8139827/, doi: 10.7554/eLife.66286.

Laturney M, Sterne GR, Scott K. Mating activates neuroendocrine pathways signaling hunger in Drosophila females. eLife. 2023 May; 12:e85117. 10.7554/eLife.85117, doi: 10.7554/eLife.85117.

Li F, Lindsey JW, Marin EC, Otto N, Dreher M, Dempsey G, Stark I, Bates AS, Pleijzier MW, Schlegel P, Nern A, Takemura Sy, Eckstein N, Yang T, Francis A, Braun A, Parekh R, Costa M, Scheffer LK, Aso Y, et al. The connectome of the adult Drosophila mushroom body provides insights into function. eLife. 2020 Dec; 9:e62576. 10.7554/eLife.62576, doi: 10.7554/eLife.62576.

Lin A, Yang R, Dorkenwald S, Matsliah A, Sterling AR, Schlegel P, Yu Sc, McKellar CE, Costa M, Eichler K, Bates AS, Eckstein N, Funke J, Jefferis GSXE, Murthy M. Network statistics of the whole-brain connectome of Drosophila. Nature. 2024 Oct; 634(8032):153–165. https://www.nature.com/articles/s41586-024-07968-y, doi: 10.1038/s41586-024-07968-y, publisher: Nature Publishing Group.

Marquis M, Wilson RI. Locomotor and olfactory responses in dopamine neurons of the Drosophila superiorlateral brain. Current Biology. 2022 Dec; 32(24):5406–5414.e5. https://www.sciencedirect.com/science/article/pii/S096098222201764X, doi: 10.1016/j.cub.2022.11.008.

Matsliah A, Sterling A, Dorkenwald S, Kuehner K, Morey R, Seung H, Murthy M. Codex: Connectome Data Explorer; 2023. doi: 10.13140/RG.2.2.35928.67844.

Matsliah A, Yu Sc, Kruk K, Bland D, Burke AT, Gager J, Hebditch J, Silverman B, Willie KP, Willie R, Sorek M, Sterling AR, Kind E, Garner D, Sancer G, Wernet MF, Kim SS, Murthy M, Seung HS. Neuronal parts list and wiring diagram for a visual system. Nature. 2024 Oct; 634(8032):166–180. https://www.nature.com/articles/s41586-024-07981-1, doi: 10.1038/s41586-024-07981-1, publisher: Nature Publishing Group.

Newman MEJ. Modularity and community structure in networks. Proceedings of the National Academy of Sciences. 2006 Jun; 103(23):8577–8582. https://www.pnas.org/doi/full/10.1073/pnas.0601602103, doi: 10.1073/pnas.0601602103, publisher: Proceedings of the National Academy of Sciences.

Newman MEJ, Girvan M. Finding and evaluating community structure in networks. Physical Review E. 2004 Feb; 69(2):026113. https://link.aps.org/doi/10.1103/PhysRevE.69.026113, doi: 10.1103/PhysRevE.69.026113, publisher: American Physical Society.

Nojima T, Rings A, Allen AM, Otto N, Verschut TA, Billeter JC, Neville MC, Goodwin SF. A sex-specific switch between visual and olfactory inputs underlies adaptive sex differences in behavior. Current Biology. 2021 Mar; 31(6):1175–1191.e6. https://www.cell.com/current-biology/abstract/S0960-9822(20)31899-6, doi: 10.1016/j.cub.2020.12.047, publisher: Elsevier.

O’Leary T, Marder E. Mapping Neural Activation onto Behavior in an Entire Animal. Science. 2014 Apr; 344(6182):372–373. https://www.science.org/doi/10.1126/science.1253853, doi: 10.1126/science.1253853.

Oram TB, Card GM. Context-dependent control of behavior in Drosophila. Current Opinion in Neurobiology. 2022 Apr; 73:102523. https://www.sciencedirect.com/science/article/pii/S0959438822000083, doi: 10.1016/j.conb.2022.02.003.

Pitman JL, Huetteroth W, Burke CJ, Krashes MJ, Lai SL, Lee T, Waddell S. A Pair of Inhibitory Neurons Are Required to Sustain Labile Memory in the *Drosophila* Mushroom Body. Current Biology. 2011 May; 21(10):855– 861. https://www.sciencedirect.com/science/article/pii/S0960982211003903, doi: 10.1016/j.cub.2011.03.069.

Prisco L, Deimel SH, Yeliseyeva H, Fiala A, Tavosanis G. The anterior paired lateral neuron normalizes odourevoked activity in the Drosophila mushroom body calyx. eLife. 2021 Dec; 10:e74172. 10.7554/eLife.74172, doi: 10.7554/eLife.74172, publisher: eLife Sciences Publications, Ltd.

Reinhard N, Fukuda A, Manoli G, Derksen E, Saito A, Möller G, Sekiguchi M, Rieger D, Helfrich-Förster C, Yoshii T, Zandawala M. Synaptic connectome of the Drosophila circadian clock. Nature Communications. 2024 Dec; 15(1):10392. https://www.nature.com/articles/s41467-024-54694-0, doi: 10.1038/s41467-024-54694-0, publisher: Nature Publishing Group.

Riva S, Ceriani MF, Risau-Gusman S, Franco DL, Circadian control of a sex-specific behaviour in Drosophila. bioRxiv; 2024. https://www.biorxiv.org/content/10.1101/2024.03.27.586991v2, doi: 10.1101/2024.03.27.586991, pages: 2024.03.27.586991 Section: New Results.

Rockwell RF, Grossfield J. Drosophila: Behavioral Cues for Oviposition. The American Midland Naturalist. 1978; 99(2):361–368. https://www.jstor.org/stable/2424813, doi: 10.2307/2424813.

Roemschied FA, Pacheco DA, Aragon MJ, Ireland EC, Li X, Thieringer K, Pang R, Murthy M. Flexible circuit mechanisms for context-dependent song sequencing. Nature. 2023 Oct; 622(7984):794–801. https://www.nature.com/articles/s41586-023-06632-1, doi: 10.1038/s41586-023-06632-1, publisher: Nature Publishing Group.

Ruta V, Datta SR, Vasconcelos ML, Freeland J, Looger LL, Axel R. A dimorphic pheromone circuit in Drosophila from sensory input to descending output. Nature. 2010 Dec; 468(7324):686–690. https://www.nature.com/articles/nature09554, doi: 10.1038/nature09554, publisher: Nature Publishing Group.

Sayin S, De Backer JF, Siju KP, Wosniack ME, Lewis LP, Frisch LM, Gansen B, Schlegel P, Edmondson-Stait A, Sharifi N, Fisher CB, Calle-Schuler SA, Lauritzen JS, Bock DD, Costa M, Jefferis GSXE, Gjorgjieva J, Grunwald Kadow IC. A Neural Circuit Arbitrates between Persistence and Withdrawal in Hungry *Drosophila*. Neuron. 2019 Nov; 104(3):544–558.e6. https://www.sciencedirect.com/science/article/pii/S0896627319306543, doi: 10.1016/j.neuron.2019.07.028.

Scheffer LK, Meinertzhagen IA. A connectome is not enough – what is still needed to understand the brain of Drosophila? Journal of Experimental Biology. 2021 Oct; 224(21):jeb242740. 10.1242/jeb.242740, doi: 10.1242/jeb.242740.

Scheffer LK, Xu CS, Januszewski M, Lu Z, Takemura Sy, Hayworth KJ, Huang GB, Shinomiya K, Maitlin-Shepard J, Berg S, Clements J, Hubbard PM, Katz WT, Umayam L, Zhao T, Ackerman D, Blakely T, Bogovic J, Dolafi T, Kainmueller D, et al. A connectome and analysis of the adult Drosophila central brain. eLife. 2020 Sep; 9:e57443. https://elifesciences.org/articles/57443, doi: 10.7554/eLife.57443.

Schlegel P, Yin Y, Bates AS, Dorkenwald S, Eichler K, Brooks P, Han DS, Gkantia M, dos Santos M, Munnelly EJ, Badalamente G, Serratosa Capdevila L, Sane VA, Fragniere AMC, Kiassat L, Pleijzier MW, Stürner T, Tamimi IFM, Dunne CR, Salgarella I, et al. Whole-brain annotation and multi-connectome cell typing of Drosophila. Nature. 2024 Oct; 634(8032):139–152. https://www.nature.com/articles/s41586-024-07686-5, doi: 10.1038/s41586-024-07686-5, publisher: Nature Publishing Group.

Schretter CE, Aso Y, Robie AA, Dreher M, Dolan MJ, Chen N, Ito M, Yang T, Parekh R, Branson KM, Rubin GM. Cell types and neuronal circuitry underlying female aggression in Drosophila. eLife. 2020 Nov; 9:e58942. 10.7554/eLife.58942, doi: 10.7554/eLife.58942, publisher: eLife Sciences Publications, Ltd.

Schwartz NU, Zhong L, Bellemer A, Tracey WD. Egg Laying Decisions in Drosophila Are Consistent with Foraging Costs of Larval Progeny. PLOS ONE. 2012 May; 7(5):e37910. https://journals.plos.org/plosone/article?id=10.1371/journal.pone.0037910, doi: 10.1371/journal.pone.0037910.

Shafer OT, Gutierrez GJ, Li K, Mildenhall A, Spira D, Marty J, Lazar AA, Fernandez MdlP. Connectomic analysis of the Drosophila lateral neuron clock cells reveals the synaptic basis of functional pacemaker classes. eLife. 2022 Jun; 11:e79139. 10.7554/eLife.79139, doi: 10.7554/eLife.79139, publisher: eLife Sciences Publications, Ltd.

Shelly TE. Defense of Oviposition Sites by Female Oriental Fruit Flies (Diptera: Tephritidae). The Florida Entomologist. 1999; 82(2):339–346. https://www.jstor.org/stable/3496587, doi: 10.2307/3496587.

Séjourné J, Plaçais PY, Aso Y, Siwanowicz I, Trannoy S, Thoma V, Tedjakumala SR, Rubin GM, Tchénio P, Ito K, Isabel G, Tanimoto H, Preat T. Mushroom body efferent neurons responsible for aversive olfactory memory retrieval in Drosophila. Nature Neuroscience. 2011 Jul; 14(7):903–910. https://www.nature.com/articles/nn.2846, doi: 10.1038/nn.2846, publisher: Nature Publishing Group.

Taisz I, Donà E, Münch D, Bailey SN, Morris BJ, Meechan KI, Stevens KM, Varela-Martínez I, Gkantia M, Schlegel P, Ribeiro C, Jefferis GSXE, Galili DS. Generating parallel representations of position and identity in the olfactory system. Cell. 2023 Jun; 186(12):2556–2573.e22. https://www.sciencedirect.com/science/article/pii/S0092867423004725, doi: 10.1016/j.cell.2023.04.038.

Vijayan V, Wang F, Wang K, Chakravorty A, Adachi A, Akhlaghpour H, Dickson BJ, Maimon G. A rise-to-threshold process for a relative-value decision. Nature. 2023 Jul; 619(7970):563–571. https://www.nature.com/articles/s41586-023-06271-6, doi: 10.1038/s41586-023-06271-6, number: 7970 Publisher: Nature Publishing Group.

Vijayan V, Wang Z, Chandra V, Chakravorty A, Li R, Sarbanes SL, Akhlaghpour H, Maimon G. An internal expectation guides Drosophila egg-laying decisions. Science Advances. 2022 Oct; 8(43):eabn3852. https://www.science.org/doi/10.1126/sciadv.abn3852, doi: 10.1126/sciadv.abn3852, publisher: American Association for the Advancement of Science.

Waddell S, Armstrong JD, Kitamoto T, Kaiser K, Quinn WG. The *amnesiac* Gene Product Is Expressed in Two Neurons in the *Drosophila* Brain that Are Critical for Memory. Cell. 2000 Nov; 103(5):805–813. https://www.sciencedirect.com/science/article/pii/S0092867400001835, doi: 10.1016/S0092-8674(00)00183-5.

Walker SR, Peña-Garcia M, Devineni AV. Connectomic analysis of taste circuits in Drosophila. bioRxiv: The Preprint Server for Biology. 2024 Sep; p. 2024.09.14.613080. doi: 10.1101/2024.09.14.613080.

Wang F, Wang K, Forknall N, Patrick C, Yang T, Parekh R, Bock D, Dickson BJ. Neural circuitry linking mating and egg laying in Drosophila females. Nature. 2020 Mar; 579(7797):101–105. http://www.nature.com/articles/s41586-020-2055-9, doi: 10.1038/s41586-020-2055-9.

Wang K, Gong J, Wang Q, Li H, Cheng Q, Liu Y, Zeng S, Wang Z. Parallel pathways convey olfactory information with opposite polarities in Drosophila. Proceedings of the National Academy of Sciences. 2014 Feb; 111(8):3164–3169. https://www.pnas.org/doi/full/10.1073/pnas.1317911111, doi: 10.1073/pnas.1317911111, publisher: Proceedings of the National Academy of Sciences.

Wang K, Wang F, Forknall N, Yang T, Patrick C, Parekh R, Dickson BJ. Neural circuit mechanisms of sexual receptivity in Drosophila females. Nature. 2021 Jan; 589(7843):577–581. https://www.nature.com/articles/s41586-020-2972-7, doi: 10.1038/s41586-020-2972-7, publisher: Nature Publishing Group.

Winding M, Pedigo BD, Barnes CL, Patsolic HG, Park Y, Kazimiers T, Fushiki A, Andrade IV, Khandelwal A, Valdes-Aleman J, Li F, Randel N, Barsotti E, Correia A, Fetter RD, Hartenstein V, Priebe CE, Vogelstein JT, Cardona A, Zlatic M. The connectome of an insect brain. Science. 2023 Mar; 379(6636):eadd9330. https://www.science.org/doi/10.1126/science.add9330, doi: 10.1126/science.add9330, publisher: American Association for the Advancement of Science.

Yang Ch, Belawat P, Hafen E, Jan LY, Jan YN. Drosophila Egg-Laying Site Selection as a System to Study Simple Decision-Making Processes. Science. 2008 Mar; 319(5870):1679–1683. https://www.science.org/doi/10.1126/science.1151842, doi: 10.1126/science.1151842, publisher: American Association for the Advancement of Science.

Yang CH, He R, Stern U. Behavioral and Circuit Basis of Sucrose Rejection by Drosophila Females in a Simple Decision-Making Task. Journal of Neuroscience. 2015 Jan; 35(4):1396–1410. https://www.jneurosci.org/content/35/4/1396, doi: 10.1523/JNEUROSCI.0992-14.2015.

Yang Ch, Rumpf S, Xiang Y, Gordon MD, Song W, Jan LY, Jan YN. Control of the Postmating Behavioral Switch in Drosophila Females by Internal Sensory Neurons. Neuron. 2009 Feb; 61(4):519–526. https://www.sciencedirect.com/science/article/pii/S0896627308010933, doi: 10.1016/j.neuron.2008.12.021.

Yu HH, Awasaki T, Schroeder MD, Long F, Yang JS, He Y, Ding P, Kao JC, Wu GYY, Peng H, Myers G, Lee T. Clonal Development and Organization of the Adult *Drosophila* Central Brain. Current Biology. 2013 Apr; 23(8):633– 643. https://www.sciencedirect.com/science/article/pii/S0960982213002649, doi: 10.1016/j.cub.2013.02.057.

Zhang L, Yu J, Guo X, Wei J, Liu T, Zhang W. Parallel Mechanosensory Pathways Direct Oviposition Decision-Making in *Drosophila*. Current Biology. 2020 Aug; 30(16):3075–3088.e4. https://www.sciencedirect.com/science/article/pii/S0960982220307636, doi: 10.1016/j.cub.2020.05.076.

Zhou C, Pan Y, Robinett CC, Meissner GW, Baker BS. Central Brain Neurons Expressing *doublesex* Regulate Female Receptivity in *Drosophila*. Neuron. 2014 Jul; 83(1):149–163. https://www.sciencedirect.com/science/article/pii/S0896627314004826, doi: 10.1016/j.neuron.2014.05.038.

